# The nucleus accumbens to ventral pallidum pathway regulates social play behavior via sex-specific mechanisms in juvenile rats

**DOI:** 10.1101/2025.09.06.674638

**Authors:** Jessica D.A. Lee, Daniela N. Anderson, Isabella C. Orsucci, Samantha M. Bowden, Alexa H. Veenema

## Abstract

Social play behavior is a rewarding behavior predominantly displayed by juveniles of various mammalian species, including humans and rats. Although the mesolimbic reward system is involved in the regulation of social play, how brain regions in this system interact to regulate social play behavior is unknown. Here, we determined the involvement of the ventral pallidum (VP) as well as inputs from the nucleus accumbens (NAc) to the VP in the regulation of social play in male and female juvenile rats. We show that acute pharmacological inactivation of the VP, via microinfusion of the GABA-A receptor agonist muscimol, decreased social play behaviors in both sexes. Next, using Gad1-iCre rats, we show that chemogenetic stimulation of NAc^GABA^ terminals in the VP decreased VP neuronal activation and decreased social play behaviors in both sexes. These findings together indicate that reduced inhibitory NAc input to the VP permits activation of the VP which facilitates the expression of social play behaviors. Lastly, we show that the equal expression of social play behavior in males and females is associated with a female-specific increase in NAc shell activation and a male-specific decrease in activation of the NAc shell neurons projecting to the VP. These sex-specific changes in NAc activity following social play exposure eliminated baseline sex differences in NAc activity. In conclusion, these findings support a model in which the sex-specific modulation of NAc inhibitory input to the VP facilitates activation of the VP that is necessary for the typical and equal expression of social play behavior in male and female juvenile rats.

## INTRODUCTION

Social play behavior is a rewarding behavior (Calcagnetti and Schechter, 1992; Ikemoto and Panksepp, 1992; Trezza et al., 2009; Achterberg et al., 2016b, 2016a, 2019) displayed by juveniles of various mammalian species, including humans and rats (Thor and Holloway, 1984a; Scott and Panksepp, 2003; Marquardt et al., 2022; Pellis et al., 2022). Social play is the earliest form of peer-to-peer social interaction (Thor and Holloway, 1984b; Pellis and Pellis, 1987), and various studies demonstrate that engagement in social play is essential for the development of social competence later in life (Bekoff, 1974; van den Berg et al., 1999). Children with autism spectrum disorder (ASD) show decreased engagement in social play behavior with their peers (Holmes and Willoughby, 2005; Buggey et al., 2011). This may be due to changes in brain regions involved in social reward processing. In support, ASD children show structural and functional changes in the mesolimbic reward system compared to their typically developing peers (Supekar et al., 2018). These changes were most prominent in the nucleus accumbens (NAc) which is a central part of the mesolimbic reward system (Salamone and Correa, 2002; Carlezon and Thomas, 2009). The NAc can be divided into two subregions, namely the NAc core and NAc shell (Heimer et al., 1991; Brog et al., 1993; O’Donnell and Grace, 1993). In male juvenile rats, pharmacological inactivation of the NAc core, but not the shell, increased social play duration (van Kerkhof et al., 2013). However, social play exposure was associated with increased neuronal activation of both the NAc core and shell in male juvenile rats (van Kerkhof et al., 2014), suggesting that both core and shell subregions are involved in the regulation of social play behavior.

The ventral pallidum (VP) is another key brain region of the mesolimbic reward system (Smith et al., 2011; Berridge and Kringelbach, 2015). We recently showed that social play exposure is associated with sex-specific changes in neuronal activation of the VP in juvenile rats (Lee et al., 2024). The VP receives major GABAergic input from the NAc (Walaas and Fonnum, 1979; Jones and Mogenson, 1980; Swerdlow et al., 1990; Root et al., 2015). This NAc GABAergic input to the VP modulates the expression of rewarding behaviors in adult rodents. For example, optogenetic stimulation of NAc terminals in the VP reduced fos mRNA expression in the VP and decreased sucrose intake in adult female rats (Chometton et al., 2020). Moreover, chemogenetic inhibition of NAc terminals in the VP decreased inhibitory postsynaptic currents in the VP and increased lever press responses and breakpoints in response to a food reward in adult male and female mice (Gallo et al., 2018). These studies suggest that NAc GABAergic inputs to the VP suppress VP activation, thereby reducing the expression of rewarding behaviors. This implies that inhibition of NAc GABAergic inputs to the VP disinhibits the VP, which is likely a necessary step for the expression of rewarding behaviors. Although the above-mentioned studies were performed in adult rodents, we propose that this pathway is similarly recruited in juvenile rats for the expression of rewarding behaviors, such as social play behavior.

Here, we determined the involvement of the VP as well as inputs from the NAc to the VP in the regulation of social play in male and female juvenile rats. We hypothesized that activation of the VP is necessary for the expression of social play in both sexes (Exp. 1), that this activation is facilitated through inhibition of the NAc to VP pathway (Exp. 2), and that this occurs via sex-specific mechanisms (Exp. 3). In Exp. 1, we determined whether pharmacological inactivation of the VP (using the GABA_A_ receptor agonist muscimol) reduces the expression of social play behaviors in male and female juvenile rats. In Exp. 2, we utilized male and female juvenile Gad1-iCre rats to determine whether chemogenetic stimulation of the NAc^GABA^ to VP pathway reduces the expression of social play behaviors. Furthermore, we determined whether chemogenetic stimulation of the NAc^GABA^ to VP pathway decreased VP neuronal activation in response to social play exposure. Lastly, in Exp. 3, we determined whether social play exposure alters the activation of NAc core and shell neurons projecting to the VP sex-specifically in juvenile rats.

## METHODS

### Animals

For Experiment 1 and 3, three-week-old male and female Wistar rats were obtained from Charles River Laboratories (Raleigh, NC). For Experiment 2, three-week-old male and female Long Evans Gad1-iCre (Sharpe et al., 2017; Gibson et al., 2018; Farrell et al., 2021, 2022) and their Cre-negative wildtype littermates were bred in-house. Stimulus rats were age-, sex-, and strain-matched but were from different litters than the experimental rats. All rats were maintained under standard laboratory conditions (12 h light/dark cycle, lights off at 14:00 h, food and water available *ad libitum*) in single-sex groups of four in standard (48 × 27 × 20 cm) or Allentown (41.2 × 30 × 23.4 × cm) rat cages unless otherwise noted. The experiments were conducted in accordance with the National Institute of Health *Guidelines for Care and Use of Laboratory Animals* and approved by the Michigan State University Institutional Animal Care and Use Committees.

### Social Play Test

Social play was assessed in juvenile (32-35 day-old) rats because social play is at its peak during this age (Panksepp, 1981; Pellis and Pellis, 1990; Paul et al., 2014). Social play testing and behavioral analysis were performed according to Veenema and Neumann (2009). Briefly, 24 hours prior to testing, experimental rats were single housed. Testing occurred during the first hour of the dark phase in which the experimental rat’s home cage was removed from the cage rack, the cage lid was removed and replaced with a Plexiglass lid. A video camera was set up above each cage to record the tests. A sex-, age-, and strain-matched stimulus rat was then placed in the cage. The social play test lasted 10 minutes, during which time the experimental rat was allowed to freely interact with the stimulus rat. Stimulus rats were striped with a permanent marker 30–60 min prior to social play testing to distinguish between the experimental and stimulus rats during later video analysis. Food and water were not available during the 10 min tests but were immediately returned upon completion of each test.

Behavior in the social play test was analyzed by a researcher blind to the sex, drug treatment, and genotype of the experimental rats using SolomonCoder software (https://solomon.andraspeter.com/). The following behaviors were scored for the experimental rats: duration of social play (the total amount of time spent in playful social interactions including nape attacks, pinning, and supine poses), duration of social investigation (the experimental rat is sniffing the anogenital and head/neck regions of the stimulus rat), duration of allogrooming (the experimental rat is grooming the stimulus rat), duration of non-social cage exploration (the experimental rat is walking, rearing, sitting, or engaging in other neutral behaviors), number of nape attacks (the experimental rat displays nose attacks or nose contacts toward the nape of the neck of the stimulus rat), number of pins (the experimental rat holds the stimulus rat on its back in a supine position), and number of supine poses (the experimental rat is pinned by the stimulus rat).

### Experimental Procedures

#### Experiment 1: Determine whether pharmacological inactivation of the VP reduces the expression of social play behavior in male and female juvenile rats

##### Bilateral VP cannulation surgery

After one week of handling, 31-day-old male and female Wistar rats underwent stereotaxic surgery. During surgery, rats were maintained under isoflurane anesthesia, 2-4% as needed (Henry Schein, Melville, NY). Guide cannulae (22 gauge, 9mm; Plastics One, Roanoke, VA) were bilaterally implanted 2 mm dorsal to the VP according to Lee et al., 2024. Coordinates for the VP from bregma were: 0.23 mm rostral to bregma, 2.4 mm lateral to the midline, and 5.8 mm ventral to the surface of the skull (Paxinos and Watson, 2007). Cannulae were implanted at an angle of 10° from the midsagittal plane to avoid damage to the sagittal sinus. Cannulae were fixed to the skull with four stainless steel screws and dental cement and closed with a dummy cannula (Plastics One, Roanoke, VA). Experimental rats were given a subcutaneous injection of Rimadyl (Covetrus 10000319; 10 mg/kg) immediately after surgery and once a day for an additional two days after surgery. Immediately after surgery, experimental rats were individually housed in standard rat cages until the end of the experiment.

##### Bilateral muscimol microinfusions followed by social play testing

Four days after surgery, at 35 days of age, experimental rats received bilateral infusions of either vehicle (0.5μL/side aCSF; pH 7.4; males n = 5; females n = 4) or the GABA_A_ receptor agonist muscimol (Sigma Aldrich, M1523; 10ng/0.5μL/side, males n = 8; females n = 7) into the VP 20 min prior to exposure to an unfamiliar stimulus rat during the 10-min social play test (described above). The infusions were given over the course of 45 s via an injector cannula (28 gauge; Plastics One, Roanoke, VA) that extended 2 mm beyond the guide cannula and was connected via polyethylene tubing to a 2 μL syringe (Hamilton Company #88400) mounted onto a microinfusion pump (GenieTouch, Kent Scientific, Torrington, CT). The injector cannula was kept in place for an additional 30 s following infusion to allow for tissue uptake before being replaced by the dummy cannula. Time of administration (Veenema et al., 2013) and drug concentration (Numan et al., 2005) were based on previous studies showing changes in social behaviors of rats upon intracerebral drug infusions.

##### Histological verification of cannula placement in the VP

At the end of the experiment, experimental rats were euthanized with CO_2_ and charcoal was injected as a marker to check proper placement of the injector cannulae. Brains were extracted, rapidly frozen in methylbutane cooled on dry ice, and stored at −80°C. Brains were sliced in 30 µm sections on the cryostat (Leica CM3050 S) and every third section was mounted directly on Superfrost Plus Slides (Fisher Scientific). Sections were stained with thionin and coverslipped with Permount mounting medium (SP15-100, Fisher Scientific, Waltham, MA). Slides were examined using light microscopy and cannula placements were mapped using The Rat Brain Atlas of Paxinos and Watson (2007). Only rats with bilateral injector cannulae tracks terminating in the VP were included in statistical analyses (Supplementary Fig 1).

#### Experiment 2A: Determine whether chemogenetic stimulation of NAc^GABA^ terminals in the VP reduces social play behaviors in male and female juvenile rats

##### Stereotaxic infusion of a cre-dependent excitatory DREADD in the NAc

At 22 days of age, Gad1-iCre Long Evans rats (males n = 6; females n = 7) and wildtype littermates (males n = 6; females n = 6) were anesthetized with isoflurane (2-4% as needed; Henry Schein, Melville, NY) and mounted on a stereotaxic frame. A 1 μL, 7000 series Hamilton syringe (Hamilton, Reno, NV) was attached to a motorized stereotaxic injector system (Stoelting, Wood Dale, IL) and bilateral injections of an excitatory DREADD-containing viral vector (0.2 μL/side; AAV-DJ-EF1a-DIO-hM3D(Gq)-mCherry; Gene Vector and Virus Core, Stanford University, CA) was directed at the NAc core and shell medial to the anterior commissure. Coordinates for the NAc from bregma were: 1.2 mm rostral to bregma, 2.7 mm lateral to the midline, and 6.7 mm ventral to the surface of the skull (Paxinos and Watson, 2007). The syringe was injected at a 10° angle from the midsagittal plane to avoid damage to the sagittal sinus. Experimental rats were given a subcutaneous injection of meloxicam (2mg/kg; Covetrus, 049759) immediately after surgery and once a day for an additional two days. Each subject was pair-housed with two novel age-, sex-, and strain-matched stimulus rats after surgery until the second stereotaxic surgery.

##### Bilateral cannulation targeting the VP and habituation to the social play test

Seven days later, at 29 days of age, all rats underwent a second stereotaxic surgery to bilaterally implant guide cannulae 2 mm above the VP, allowing for local infusion of the synthetic ligand clozapine-n-oxide (CNO) or 0.9% sterile saline. Bilateral cannulation procedures were the same as described in Experiment 1. Post-operative monitoring was the same as the first stereotaxic surgery, with the exception that all rats were also given a subcutaneous injection of Enrofloxacin (22.7mg/kg; Covetrus, 074743) immediately after surgery. Following surgery, each experimental rat was rehoused with the same two age- and sex-, and strain-matched stimulus rats. One day later, at 30 days of age, experimental rats were socially isolated by removing the two stimulus rats from the homecage. The two stimulus rats were placed into a novel homecage and remained pair-housed for the duration of the behavioral experiment. At 31 days of age, experimental rats were habituated to the infusion procedure and the 10-min social play test.

##### Bilateral CNO microinfusions followed by social play testing

At 32 and 33 days of age, experimental rats received, in counterbalanced order, bilateral infusions of either saline (0.9% sterile saline; Medline, 533-JB1301P) or CNO (1mM; dissolved in 0.2M sterile PBS with 10% (2-Hydroxypropyl)-β-cyclodextrin) into the VP 20 min prior to exposure to a familiar stimulus rat in the social play test (described above). The microinfusion procedure was the same as described in Experiment 1.

##### Histological verification of DREADD transduction in the NAc and of cannula placement in the VP

At 34 days of age, experimental rats were deeply anesthetized with isoflurane before being transcardially perfused with 0.9% saline followed by 4% paraformaldehyde in 0.1M phosphate buffer (pH: 9.5) euthanized via transcardial perfusions. Brains were then extracted, post-fixed for 24 hrs in 12% sucrose in 4% paraformaldehyde, rapidly frozen in methylbutane, and stored at 80°C. Brains were cryocut (Leica CM3050 S) into four series of 30 μm coronal sections containing the NAc (corresponding to distances +3.24 mm to +1.08 mm from bregma; Paxinos and Watson, 2007) and VP (corresponding to distances +0.96 mm to −0.24 mm from bregma; Paxinos and Watson, 2007). All four series were put into a cryoprotectant solution (0.05 mol L−1 sodium phosphate buffer, 30% ethylene glycol, 20% glycerol) and stored at −20°C until further histological processing.

For Gad1-iCre experimental rats, one series of the NAc was processed using fluorescence mCherry immunohistochemistry to determine DREADDs transduction and co-stained with a fluorescent Nissl to identify cytoarchitectonic borders of brain regions. Briefly, tissue sections were thoroughly rinsed in tris buffered saline (TBS; pH: 7.4) and incubated for 24 hrs at 4°C in a blocking solution (TBS with 0.3% Triton X-100 and 2% normal donkey serum; 017-000-121; Jackson ImmunoResearch, West Grove, PA) with the primary antibody anti-mCherry raised in chicken (1:2000 concentration; AB205402, ABCam). Afterwards, tissue sections were rinsed in TBS and incubated for 1 hr in the blocking solution containing the secondary antibody Alexa Fluor 594 anti-chicken raised in donkey (1:500 concentration; 703-585-155, Jackson ImmunoResearch). After the secondary antibody incubation, tissue sections were rinsed in TBS and stained with a fluorescent Nissl (1:500, 1 hr; NeuroTraceTM; N21479, 435/455nm, Thermo Fisher Scientific) at room temperature. Sections were then mounted onto gelatin-coated slides, air-dried, and coverslipped with Vectashield hardset antifade mounting medium with a DAPI counterstain (H-1500-10, Vector Laboratories) and stored at 4°C. DREADD expression was visualized with a 4x objective on a Keyence BZ-X700E/BZ-X710 fluorescent microscope and associated BZ-H3AE software (Keyence Corporation of America). Gad1-iCre rats showing bilateral DREADDs transduction in the NAc core and shell (Supplementary Fig 2a-e) were included in the final analysis.

For experimental wildtype rats and Gad1-iCre rats receiving DREADDs infusion into the NAc, one series of the VP was stained using a fluorescent Nissl (as described above) to determine cannula placements using the Rat Brain Atlas (Paxinos and Watson, 2007). Wildtype and Gad1-iCre rats with bilateral cannula tracks terminating in the VP (Supplementary Fig 2f, 2g) were included in behavioral scoring and statistical analysis.

#### Experiment 2B. Determine whether chemogenetic stimulation of NAc^GABA^ terminals in the VP decreases VP neuronal activation in response to social play exposure

##### Unilateral CNO and contralateral saline microinfusions followed by social play testing

At 34 days of age, experimental wildtype and Gad1-iCre rats from Experiment 2A received CNO into the VP of one hemisphere and 0.9% sterile saline into the VP of the contralateral hemisphere 20 min prior to exposure to the social play test. Thirty minutes after the start of the 10-min social play test, experimental rats were deeply anesthetized with isoflurane before being transcardially perfused with 0.9% saline followed by 4% paraformaldehyde in 0.1M phosphate buffer (pH: 9.5). This time course was chosen because stimulus-induced c-fos mRNA expression is at its peak 30 min after stimulation (Morgan and Curran, 1991). Brains were then extracted and post-fixed for 24 hrs in 12% sucrose in 4% paraformaldehyde, rapidly frozen in methylbutane and stored at 80°C.

##### Histological quantification of fos-positive cells in the VP

Brains were cryocut in four series containing the VP as described in Experiment 2A. The second series of VP was processed for fluorescent *in situ* hybridization to visualize *fos* mRNA-expressing cells in the VP in a subset of experimental rats from Experiment 2A (wildtype: males n = 2, females n = 3; Gad1-iCre: males n = 2, females n = 3). Blocked tissue sections containing only the VP were mounted onto separate slides (Superfrost Plus; Fisher Scientific). RNAScopeTM Multiplex Fluorescent Reagent V2 Kits (323100, Advanced Bell Diagnostics) and probes to detect *fos* mRNA were used according to user manual from the supplier (Document Number 323100-USM, Advanced Cell Diagnostics). Briefly, tissue sections were washed in phosphate buffer solution (pH: 7.6), dried at 60°C (30 min), then post-fixed in 4% paraformaldehyde (15 min) followed by dehydration in an ethanol series. Following hydrogen peroxide incubation (10 min) and target retrieval in a steamer at 99°C (5 min), tissue was then treated with protease III (30 min; 322340, Advanced Cell Diagnostics) at room temperature. The fos-C1 (403591, Advanced Cell Diagnostics) probe was then hybridized in a HybEZTM oven (2 hrs; Advanced Cell Diagnostics) at 40°C. After probe hybridization, tissue sections were incubated with amplifier probes (AMP1, 40°C, 30 min; AMP2, 40°C, 30 min; AMP3, 40°C, 15 min). *fos* mRNA was tagged to the fluorophore fluorescein (1:1500; NEL741E001KT, Akoya Biosciences, 40°C, 30 min). Slides were then rinsed in TBS and stained with a fluorescent Nissl (1:500, 1 hr; NeuroTraceTM; N21479, 435/455nm, Thermo Fisher Scientific). Slides were then coverslipped with Vectashield hardset antifade mounting medium with a DAPI counterstain (H-1500-10, Vector Laboratories) and stored at 4°C. All images were acquired with a 40X objective on a Keyence BZ-X700E/BZ-X710 fluorescent microscope and associated BZ-H3AE software (Keyence Corporation of America). Cells were counted as *fos*+ if they had five or more puncta (Farrell et al., 2021). In each image, the total number of *fos*+ cells were counted by the experimenter blind to sex, genotype, and drug treatment of each hemisphere.

#### Experiment 3: Determine whether social play exposure alters activation of NAc core and shell neurons projecting to the VP sex-specifically in juvenile rats

##### Stereotaxic microinfusion of CtB into the VP and social play exposure

At 25 days of age, male (n = 12) and female (n = 10) Wistar rats were anesthetized with isoflurane (2-4% as needed; Henry Schein, Melville, NY) and mounted on a stereotaxic frame. A 1 μL, 7000 series Hamilton syringe (Hamilton, Reno, NV) was attached to a motorized stereotaxic injector system (Stoelting, Wood Dale, IL) and a 0.2 μL unilateral injection of the retrograde tracer cholera toxin-B (CtB) conjugated to a fluorescent fluorophore (Alexa Fluor 594, Molecular Probes, dissolved in 0.1M PBS, resulting in 1% CtB solution) was directed to the left hemisphere of the VP at a rate of 0.1 μL/min using the following coordinates: 0.23 mm rostral to bregma, 2.5 mm lateral to the midline, and 7.7 mm ventral to the surface of the skull (Paxinos and Watson, 2007). Injections were made under an angle of 10° from the midsagittal plane to avoid damage to the sagittal sinus. The needle was left in place for 10 min following the injection to allow time for tissue uptake of the tracer. Experimental rats were given a subcutaneous injection of meloxicam (2mg/kg; Covetrus, 049759) immediately after surgery and once a day for an additional two days. Immediately after surgery, experimental rats were pair-housed with an age-, sex-, and strain-matched rat and remained pair-housed until behavioral testing. All rats remained undisturbed for seven days to allow the tracer to be taken up by axon terminals and transported back to the cell bodies of origin.

One week later, at 33 days of age, experimental rats were divided into “No Social Play” (males n = 6; females n = 4) and “Social Play” (males n = 6; females n = 6) conditions. All experimental rats were single-housed in new cages, and the pair-housed cages of the rats in the “Social Play” condition were kept and used as the social play testing environment. The following day, experimental rats in the “Social Play” condition were rejoined with their previous cagemate in their original pair-housed cage for the 10-min social play test. One of the rats in each pair was striped with a permanent marker 30-60 min prior to social play testing to distinguish the two rats during later video analysis. After the 10 min social play test, rats were returned to their single-housed cages. Thirty minutes after the start of the 10 min social play test, rats in the “Social Play” condition were euthanized via transcardial perfusions. The cages of the rats in the “No Social Play” condition were removed from the cage rack and placed in the testing area, similar to the cages for rats in the “Social Play” condition. However, rats in the “No Social Play” condition remained single-housed until euthanasia via transcardial perfusions 40 min later. Perfusion and post-fixation procedures were performed as described in Experiment 2A.

##### Histological quantification of fos-positive NAc core and shell neurons projecting to the VP

Brains were cryocut in four series of 30 μm coronal sections containing the NAc and VP as described in Experiment 2A. One series from the NAc and VP was stained with a fluorescent Nissl (1:500, 1 hr; NeuroTraceTM; N21479, 435/455nm, ThermoFisherScientific) to assess CtB infusion sites using a Keyence BZ-X700E/BZ-X710 fluorescent microscope. CtB infusion sites were mapped using The Rat Brain Atlas of Paxinos and Watson (2007). Only rats with CtB infusion sites restricted to the VP were included in the NAc fos and CtB analyses (Fig. 9C). The second series of the NAc was processed for fluorescent *in situ* hybridization to visualize *fos* mRNA-expressing neurons in the NAc using the same procedure as described in Experiment 2B. All images were acquired with a 40x objective on a Keyence BZ-X700E/BZ-X710 fluorescent microscope and associated BZ-H3AE software (Keyence Corporation of America). Images were taken at three sampling locations across the NAc core and shell. Specifically, the following anterior-posterior distances were imaged: +1.68 mm, +1.32 mm and +1.08 mm from bregma according to Paxinos and Watson (2007). At each anterior-posterior location, a dorsal and ventral image was taken (See Supplementary Fig 4 for the imaging plan). Cells were counted as *fos*+ if they had five or more *fos* puncta and CtB+ if they had one or more CtB puncta. The number of *fos*+ cells, CtB+ cells, *fos*+ cells that co-express CtB, the percent of *fos*+ cells that co-expressed CtB [(# of double-labeled cells/total number of *fos*+ cells)*100], and the percent of CtB+ cells that co-express *fos* [(# of double-labeled cells/total number of CtB+ cells)*100] were quantified.

### Statistical analysis

For Experiment 1, a two-way analysis of variances (ANOVA) was used to determine the effects of drug treatment (muscimol vs vehicle; between-subjects factor) and sex on behaviors during the social play test. For Experiment 2A, a mixed-effects ANOVA was used to determine the effects of sex and drug treatment (saline vs CNO; within-subjects factor) on behaviors during the social play test. Separate ANOVAs were used for wildtype and for Gad1-iCre rats. For Experiment 2B, a paired sample t-test was used to determine the effects of drug treatment (saline vs CNO) on the number of *fos*+ cells in the VP. Separate t-tests were used for wildtype and Gad1-iCre rats. For Experiment 3, an independent sample t-test was used to analyze the effect of sex on behaviors during the social play test. A mixed-effects ANOVA was used to assess the effects of sex, social play exposure (social play vs no social play; between-subjects factor), and NAc sampling location (anterior vs posterior; within-subjects factor) on the number of *fos*+ cells, CtB+ cells, *fos*+ cells that co-express CtB, the percent of *fos*+ cells that co-expressed CtB, and the percent of CtB+ cells that co-express *fos*. Because there was no main effect of, or interaction with, sampling location, data was collapsed across sampling locations. Pearson correlation was used to determine whether there was an association between the number of *fos*+ cells, CtB+ cells, *fos*+ cells that co-express CtB, the percent of *fos*+ cells that co-expressed CtB, and the percent of CtB+ cells that co-express *fos* in the NAc core and shell with the time spent engaging in social play behavior. When significant interactions were found, Bonferroni *post hoc* tests were conducted to clarify the effects. All data were analyzed using GraphPad Prism 10 or IBM SPSS 28, and statistical significance was set at p < 0.05. Partial eta squared (η_p_^2^) for all mixed-effects models and Cohen’s d (*d*) effect sizes for the t-test were manually computed when significant main effects or interactions were found.

## RESULTS

### Experiment 1: Pharmacological inactivation of the VP via bilateral microinfusions of muscimol reduced social play behaviors in male and female juvenile rats

Muscimol-treated rats displayed less social play, fewer nape attacks, and fewer pins compared to vehicle-treated rats (See Table 1 for statistical details; Fig 1a-c). In contrast, muscimol-treated rats showed a similar duration of social investigation (Fig 1d) and allogrooming (Fig 1e) compared to vehicle-treated rats. This indicates that the effects of muscimol were not generalized to social behaviors but rather, were specific to social play. Moreover, muscimol-treated rats showed more non-social cage exploration than vehicle-treated rats (Fig 1f). This indicates that the muscimol-induced decrease in social play behaviors is not due to a decrease in overall locomotion. Finally, there were no main or interaction effects of sex on any of the other behaviors analyzed.

**Figure 1.**
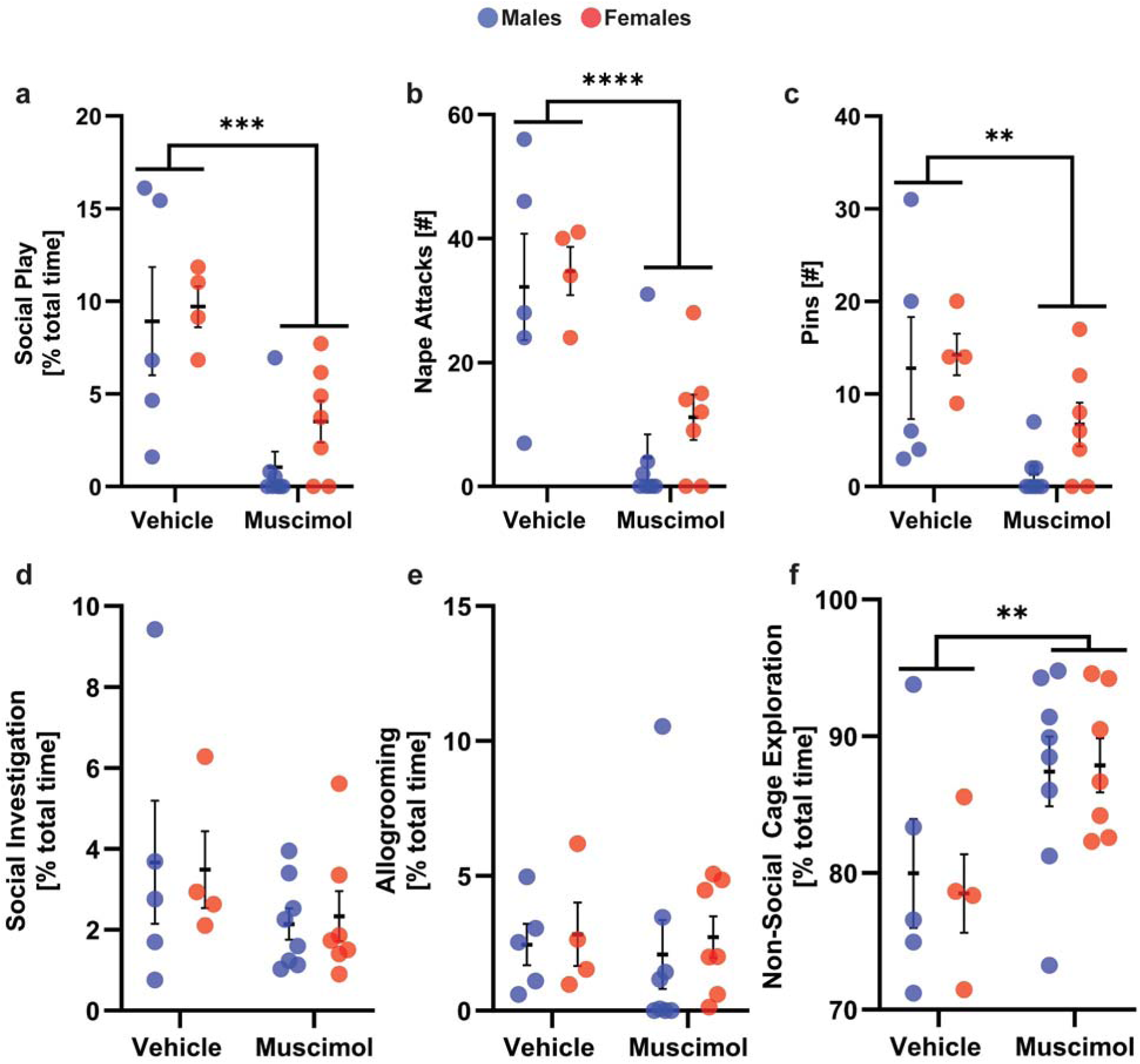
Bilateral infusions of muscimol into the ventral pallidum reduced the expression of social play behaviors in male and female juvenile rats. Bilateral ventral pallidal infusions of the GABA_A_ receptor agonist muscimol decreased the duration of social play (a), the number of nape attacks (b), and the number of pins (c) in males and females. Muscimol treatment did not alter the duration of social investigation (d) or allogrooming (e) but increased the duration of non-social cage exploration (f). Black bars indicate mean ± SEM; \**p* < 0.05, \*\**p* < 0.01, \*\*\*\**p* < 0.0001, treatment effects, two-way ANOVA.

**Table 1.**
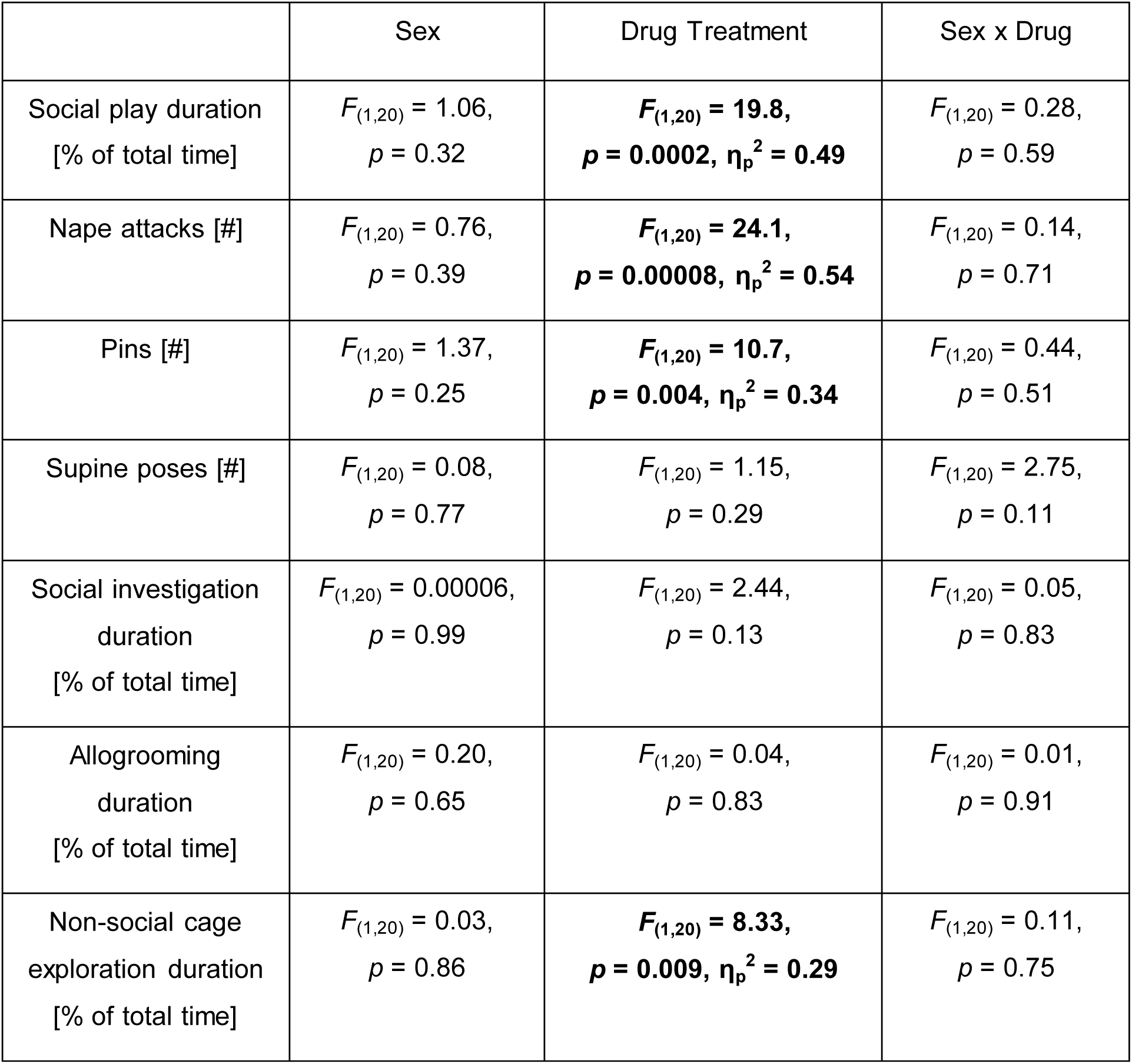
Experiment 1: Two-way ANOVA statistics for behaviors of muscimol-treated and vehicle-treated juvenile rats exposed to the social play test. Significant effects are indicated in **bold**.

### Experiment 2A: Chemogenetic stimulation of the NAc^GABA^ to VP pathway reduced social play behaviors in male and female juvenile rats

There was a main effect of drug in Gad1-iCre rats, in which CNO infusion in the VP decreased social play duration, as well as the number of nape attacks, pins, and supine poses compared to saline administration, regardless of sex (see Table 2 for statistical details; Fig 2a-c). CNO treatment did not alter the duration of social investigation or allogrooming (Fig 2d-e). This indicates that the effects of CNO were not generalized to social behaviors but rather, were specific to social play. Additionally, CNO treatment increased non-social cage exploration (Fig 2f), suggesting that the CNO-induced decrease in social play behaviors was not due to an overall decrease in locomotion.

**Figure 2.**
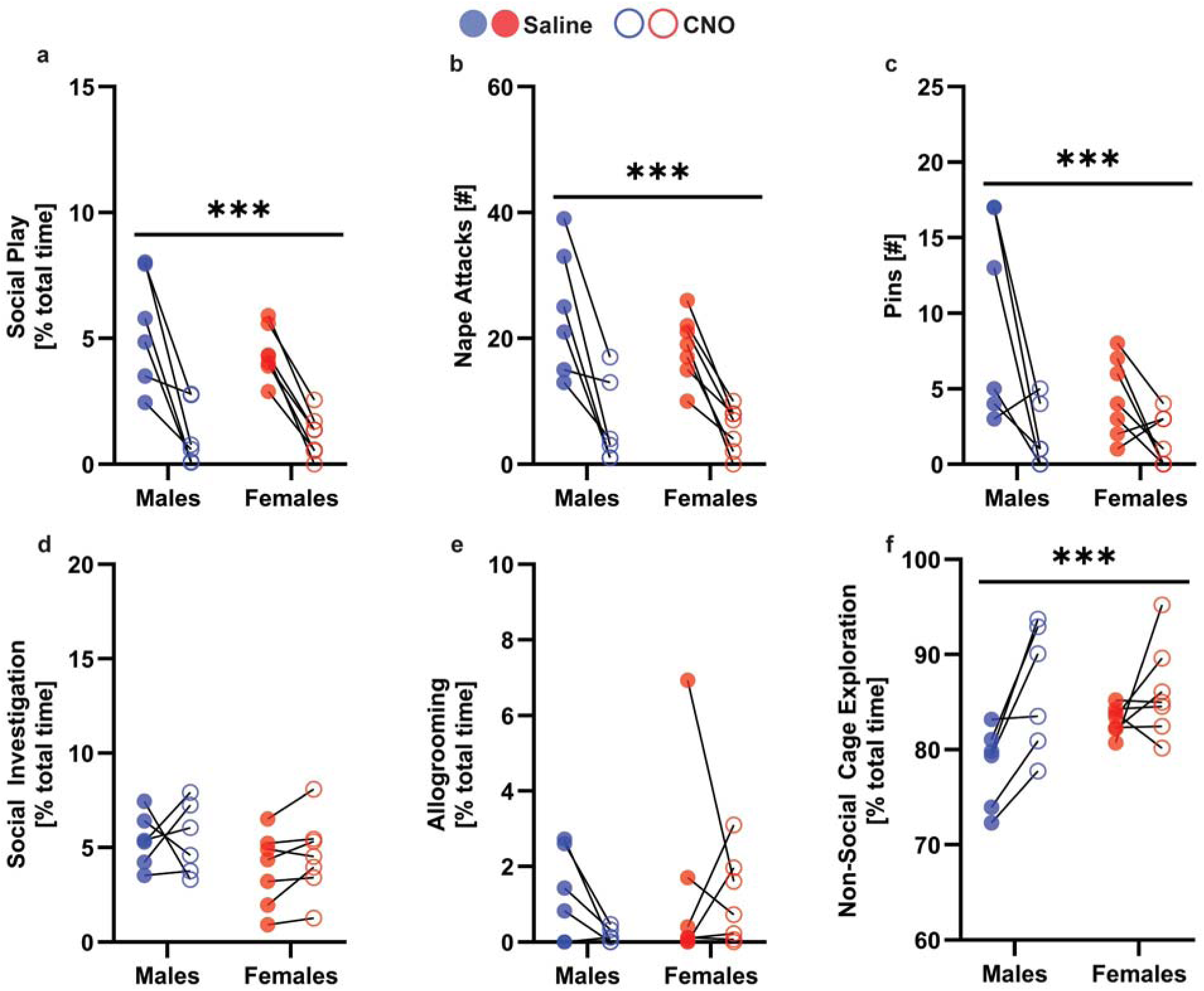
Chemogenetic stimulation of NAc^GABA^ terminals in the VP reduced social play behaviors in male and female juvenile rats. Gad1-iCre rats showed a shorter duration of social play (a), fewer nape attacks (b), and fewer pins (c) when treated with CNO compared to when they were treated with saline. Gad1-iCre rats showed similar durations of social investigation (d) and allogrooming (c) following saline and CNO treatments. Gad1-iCre rats showed a higher duration of non-social cage exploration (g) when treated with CNO compared to when they were treated with saline. ***p < 0.0001, treatment effect, mixed effects ANOVA.

**Table 2.** Experiment 2A: Two-way ANOVA statistics for behaviors of CNO-treated and saline-treated Gad1-iCre rats exposed to the social play test. Significant effects are indicated in **bold**.

|  | Sex | Drug Treatment | Sex x Drug |
| --- | --- | --- | --- |
| Social play duration<br>[% of total time] | $F_{(1,11)} = 0.68,$<br>$p = 0.42$ | **$F_{(1,11)} = 58.67,$**<br>**$p = 0.0001, \eta_p^2 = 0.84$** | $F_{(1,11)} = 1.01,$<br>$p = 0.33$ |
| Nape attacks [#] | $F_{(1,11)} = 1.23,$<br>$p = 0.29$ | **$F_{(1,11)} = 49.05,$**<br>**$p = 0.0001, \eta_p^2 = 0.92$** | $F_{(1,11)} = 1.21,$<br>$p = 0.29$ |
| Pins [#] | $F_{(1,11)} = 4.02,$<br>$p = 0.07$ | **$F_{(1,11)} = 13.31,$**<br>**$p = 0.003, \eta_p^2 = 0.54$** | $F_{(1,11)} = 2.98,$<br>$p = 0.11$ |
| Supine poses [#] | $F_{(1,11)} = 1.70,$<br>$p = 0.21$ | **$F_{(1,11)} = 7.11,$**<br>**$p = 0.02, \eta_p^2 = 0.39$** | $F_{(1,11)} = 1.70,$<br>$p = 0.21$ |
| Social investigation<br>duration<br>[% of total time] | $F_{(1,11)} = 1.83,$<br>$p = 0.20$ | $F_{(1,11)} = 0.54,$<br>$p = 0.47$ | $F_{(1,11)} = 0.32,$<br>$p = 0.58$ |
| Allogrooming duration<br>[% of total time] | $F_{(1,11)} = 0.55,$<br>$p = 0.47$ | $F_{(1,11)} = 1.30,$<br>$p = 0.27$ | $F_{(1,11)} = 0.56,$<br>$p = 0.46$ |
| Non-social cage<br>exploration duration<br>[% of total time] | $F_{(1,11)} = 1.19,$<br>$p = 0.29$ | **$F_{(1,11)} = 13.26,$**<br>**$p = 0.003, \eta_p^2 = 0.55$** | $F_{(1,11)} = 2.84,$<br>$p = 0.12$ |

CNO infusions did not alter the expression of any behaviors in the social play test compared to saline infusions in male and female wildtype rats (Supplementary Fig 3; Supplementary Table 1), demonstrating that CNO did not alter social play behaviors in the absence of DREADDs transduction. There were no main or interaction effects of sex on the expression of social play behaviors in Gad1-iCre (Table 2) and wildtype rats (Supplementary Table 1). However, there was a main effect of sex on the duration of non-social cage exploration (Supplementary Fig 3f), with females showing more non-social cage exploration than males.

### Experiment 2B: Chemogenetic stimulation of NAc^GABA^ terminals in the VP reduces VP neuronal activation in male and female juvenile rats

Gad1-iCre rats showed hM3Dq-mCherry-positive cells in the NAc core and shell (Supplementary Fig 2b, 2e) and hM3Dq-mCherry-positive fibers in the VP (Supplementary Fig 2d). As expected, WT rats did not show hM3Dq-mCherry expression in either the NAc or VP (not shown).

The number of *fos*+ cells was similar between the saline-infused and CNO-infused VP hemispheres in WT rats (*t_(4)_* = 0.89, *p* = 0.42; Fig 3c). However, there was a main effect of drug on the number of *fos*+ cells in Gad1-iCre rats, such that the CNO-infused VP hemisphere had fewer *fos*+ cells compared to the saline-infused VP hemisphere (*t_(4)_* = 2.96, *p* = 0.04, *d* = 1.34; Fig 3d). Using *fos* mRNA as an indirect marker of neuronal activity, these results confirm suppression of VP neuronal activity by CNO-induced activation of hM3Dq receptors located on NAc^GABA^ terminals in the VP.

**Figure 3.**
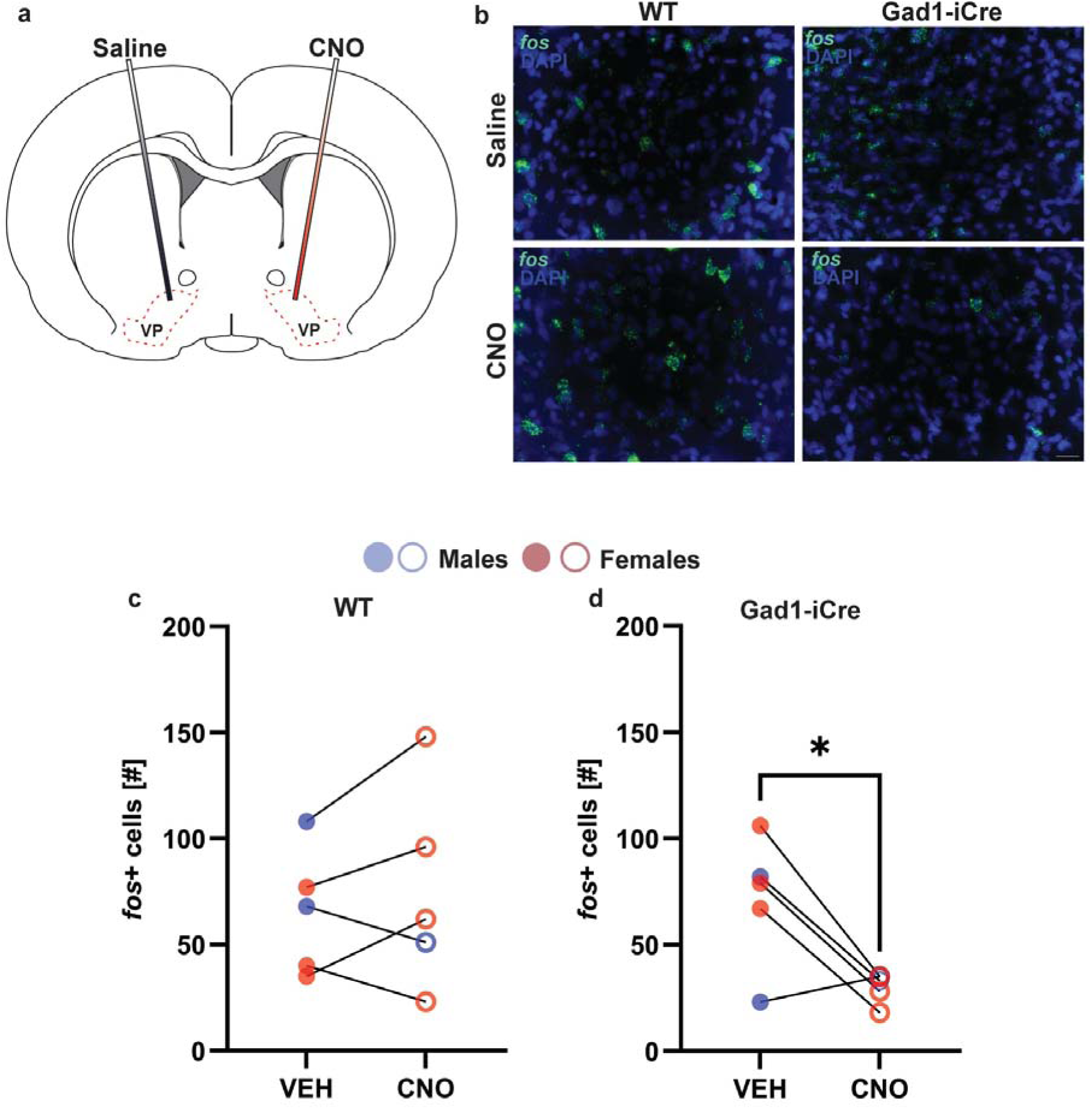
Chemogenetic stimulation of NAc^GABA^ terminals in the VP reduces VP neuronal activation in male and female juvenile rats. (a) Schematic illustration of saline and CNO microinfusion in VP hemispheres of wildtype (WT) and Gad1-iCre rats with saline (VEH) infused in one VP hemisphere and CNO infused in the contralateral VP hemisphere. (b) Representative photomicrographs showing *fos* mRNA positive (*fos*+) cells (green) in the VP following microinfusions of either saline or CNO in wildtype and Gad1-iCre rats. (c) Widltype rats displayed a similar number of *fos*+ cells in the VP hemisphere infused with saline compared to the contralateral VP hemisphere infused with CNO. (d) Gad1-iCre rats displayed a lower number of *fos*+ cells in the VP hemisphere infused with CNO compared to the VP hemisphere infused with saline. *p < 0.05, paired samples t-test.

### Experiment 3: NAc neurons projecting to the VP are sex-specifically activated in juvenile rats upon social play exposure

#### Males and females show a similar number of NAc neurons projecting to the VP

The centers for CtB infusions in the VP were similar between males and females in the “No Social Play” and “Social Play” groups and were localized between +0.48 mm and +0.24 mm from bregma (Paxinos and Watson, 2007; Supplementary Fig 5a-c). Moreover, males and females in the “No Social Play” and “Social Play” groups showed a similar number of CtB+ cells in both the NAc core and NAc shell (Supplementary Fig 5d-e; Supplementary Table 2).

#### Social play exposure is associated with increased activation of the NAc core in both sexes

Males and females in the “Social Play” group displayed a similar duration of social play (Fig 4a) and showed an increase in the number of *fos*+ cells in the NAc core compared to the “No Social Play” group (Main effect of social play condition; Table 3; Fig 4b-c). There was no significant correlation between social play duration and the number of *fos*+ cells in the NAc core in either sex (Supplementary Fig 7a).

**Figure 4.**
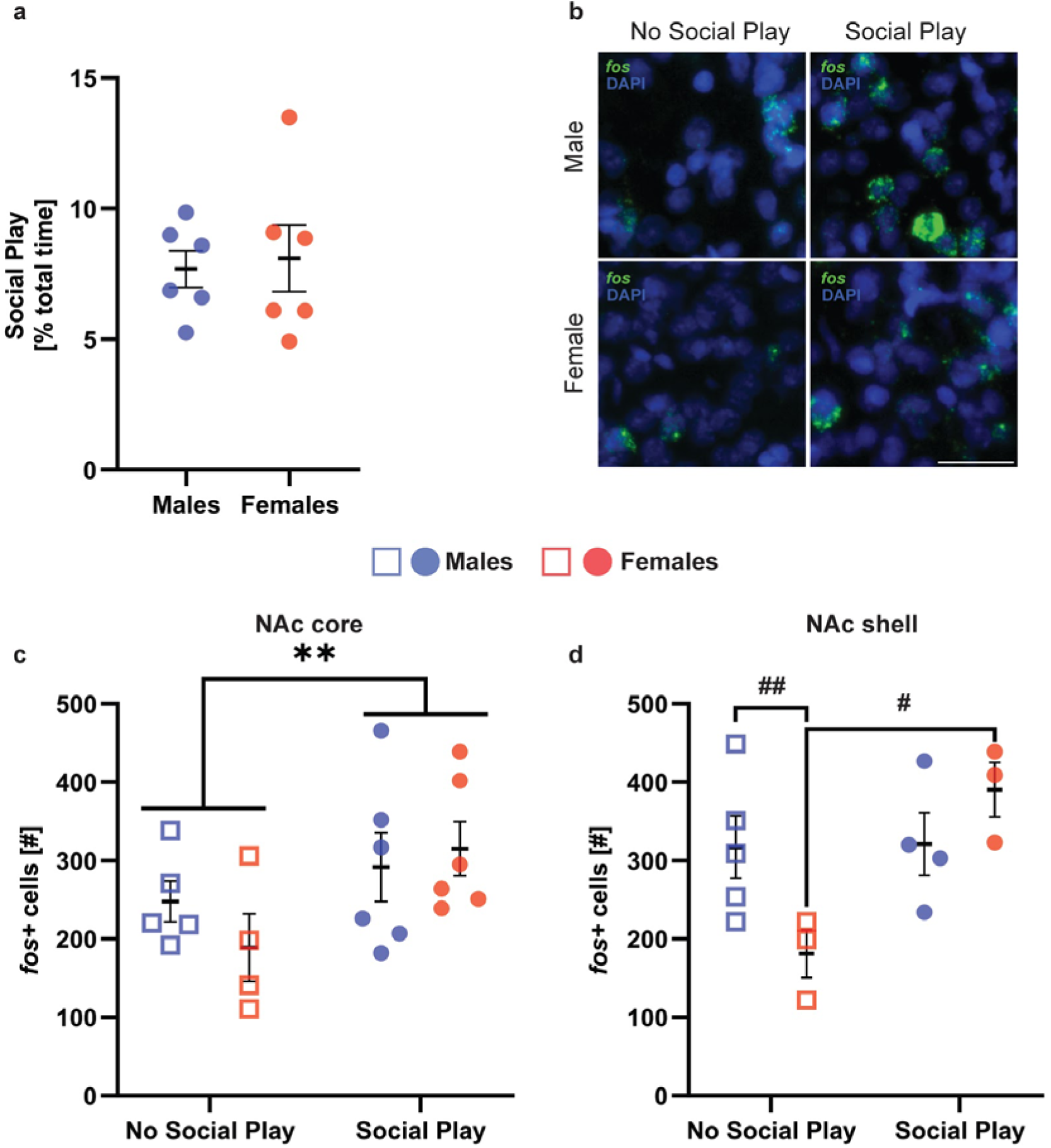
Social play exposure is associated with increased activation of the NAc core in both sexes and increased activation of the NAc shell in females only. (a) Male and female juvenile rats in the “Social Play” group showed similar durations of social play. (b) Example photomicrographs showing *fos* mRNA positive (*fos*+) cells (green) in the NAc core. (c) Males and females in the “Social Play” group showed a greater number of *fos*+ cells compared to the “No Social Play” group. (d) Males in the “No Social Play” group showed a greater number of *fos*+ cells in the NAc shell compared to females in the “No Social Play” group. In addition, females in the “Social Play” group showed a greater number of *fos*+ cells in the NAc shell than females in the “No Social Play” group. Black bars indicate mean ± SEM; \*\**p* < 0.001, main effect, two-way ANOVA; #*p* < 0.05, ##*p* < 0.001, Bonferroni *post hoc* tests. Scale bar = 15 μm.

**Table 3.** Experiment 3: Two-way ANOVA statistics for the number of *fos*+ cells in the NAc core and shell in the “No Social Play” and “Social Play” groups. Significant effects are indicated in **bold**.

|  | Sex | Social Play Condition | Sex x Social Play Condition |
| --- | --- | --- | --- |
| NAc core *fos*+ cells [#] | $F_{(1,17)} = 0.21$,<br>$p = 0.65$ | **$F_{(1,17)} = 4.83$,**<br>**$p = 0.04$, $\eta_p^2 = 0.22$** | $F_{(1,17)} = 1.13$,<br>$p = 0.30$ |
| NAc shell *fos*+ cells [#] | $F_{(1,11)} = 0.68$,<br>$p = 0.42$ | **$F_{(1,11)} = 7.06$,**<br>**$p = 0.02$, $\eta_p^2 = 0.39$** | **$F_{(1,11)} = 6.54$,**<br>**$p = 0.02$, $\eta_p^2 = 0.37$** |

#### Social play exposure is associated with increased activation of the NAc shell in females only

There were main effects of social play condition and sex x social play condition on the number of *fos*+ cells in the NAc shell (Table 3). Bonferroni *post hoc* testing revealed that the sex difference in the number of *fos*+ cells in the NAc shell (males > females) was eliminated following social play exposure due to an increase in the number of *fos*+ cells in the NAc shell in females only (Fig. 4d). There was no significant correlation between social play duration and the number of *fos*+ cells in the NAc shell in either sex (Supplementary Fig 8a).

#### Social play exposure is associated with decreased activation of NAc core and shell neurons projecting to the VP in males only

There were significant sex x social play condition effects on the number of CtB+ cells that co-expressed *fos* and the proportion of *fos*+ cells that co-expressed CtB in the NAc core and shell (Table 4). Bonferroni *post hoc* testing revealed that the sex difference in the number of CtB+ cells that co-expressed *fos* in the NAc core and shell (males > females; Fig 5b & c) was eliminated by social play exposure due to females showing a trend towards more CtB+ cells co-expressing *fos* in the NAc core (Fig 5b) and due to males showing significant fewer CtB+ cells that co-expressed *fos* in the NAc shell (Fig 5c). Furthermore, social play exposure in males reduced the proportion of *fos*+ cells that co-expressed CtB in the NAc core and shell (Fig 5d & e), thereby eliminating the baseline sex difference in the proportion of *fos*+ cells that co-expressed CtB in the NAc core (Fig 5d). There was no significant correlation between social play duration and the number of CtB+ cells that co-expressed *fos* or the proportion of *fos*+ cells that co-expressed CtB in males or females in the NAc core and shell (Supplementary Fig 7, 8).

**Figure 5.**
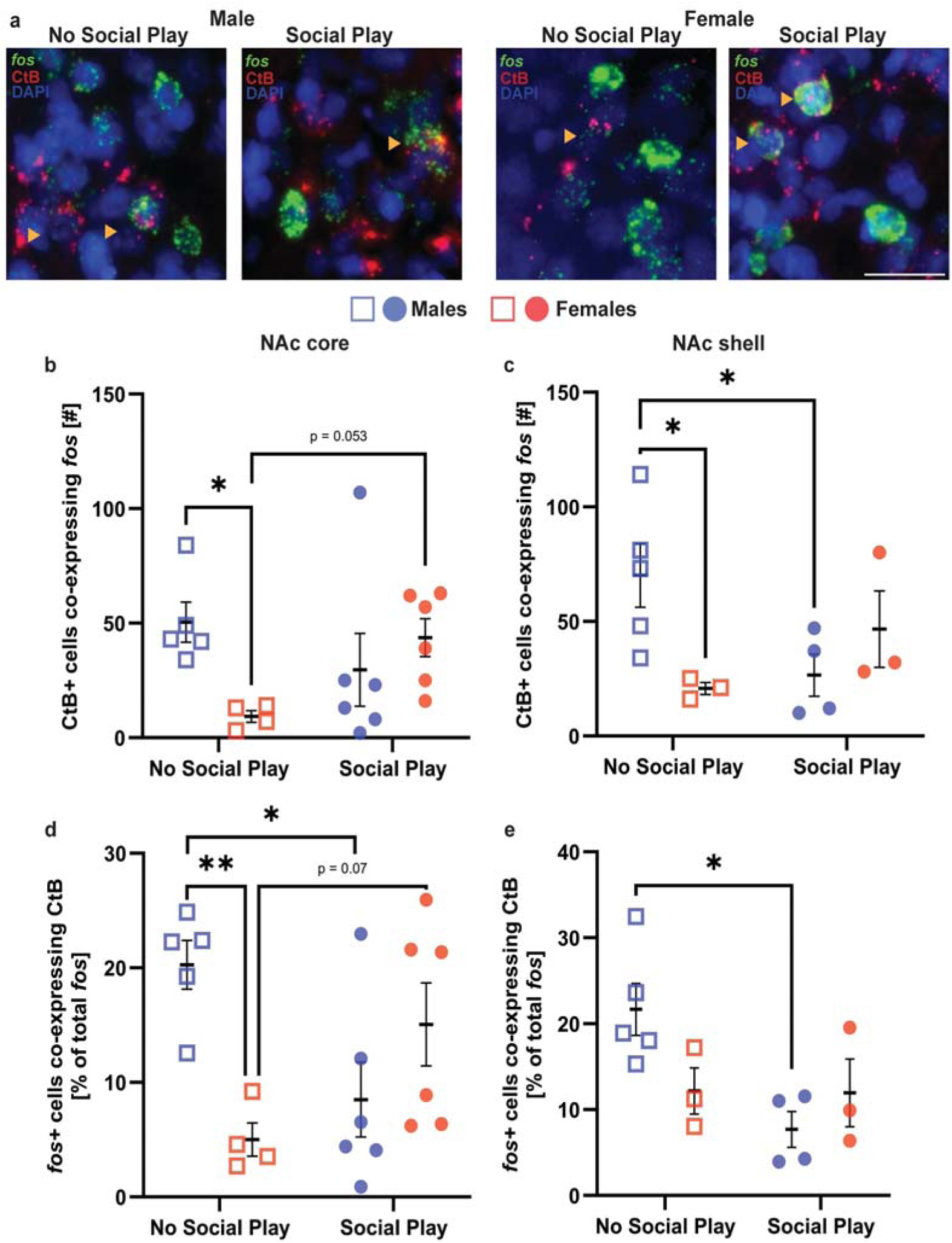
Social play exposure is associated with decreased activation of NAc core and shell neurons projecting to the VP in males, but not in females. (a) Example photomicrograph showing *fos* mRNA positive (*fos*+) cells (green) and CtB+ cells (red). Yellow arrowheads indicate *fos*+ cells co-expressing CtB. (b) The number of VP-projecting NAc core cells that co-expressed *fos* was greater in males compared to females in the “No Social Play” group but was similar between males and females in the “Social Play” group. (c) The number of VP-projecting NAc shell cells that co-expressed *fos* was greater in males in the “No Social Play” group compared to females in the “No Social Play” group and compared to males in the “Social Play” group. (d) The proportion of *fos*+ NAc core cells projecting to the VP was greater in males in the “No Social Play” group compared to females in the “No Social Play” group and compared to males in the “Social Play” group. (e) The proportion of *fos*+ NAc shell cells projecting to the VP was greater in males in the “No Social Play” group compared to males in the “Social Play” group. Black bars indicate mean ± SEM; *p < 0.05, **p < 0.001, Bonferroni *post hoc* tests following two-way ANOVA.

**Table 4.** Experiment 3: Two-way ANOVA statistics for the number of and proportion of *fos*+ cells co-expressing CtB in the NAc core and shell in the “No Social Play” and “Social Play” groups. Significant effects are indicated in **bold**.

|  | Sex | Social Play Condition | Sex x Social Play Condition |
| --- | --- | --- | --- |
| NAc core *fos*+ cells co-expressing CtB [#] | $F_{(1,17)} = 1.43$,<br>$p = 0.24$ | $F_{(1,17)} = 0.36$,<br>$p = 0.55$ | **$F_{(1,17)} = 5.90$,**<br>**$p = 0.02$, $\eta_p^2 = 0.25$** |
| NAc shell *fos*+ cells co-expressing CtB [#] | $F_{(1,11)} = 1.27$,<br>$p = 0.28$ | $F_{(1,11)} = 0.45$,<br>$p = 0.51$ | **$F_{(1,11)} = 7.23$,**<br>**$p = 0.02$, $\eta_p^2 = 0.39$** |
| NAc core *fos*+ cells co-expressing CtB [% total *fos*] | $F_{(1,17)} = 1.97$,<br>$p = 0.17$ | $F_{(1,17)} = 0.07$,<br>$p = 0.78$ | **$F_{(1,17)} = 12.49$,**<br>**$p = 0.002$, $\eta_p^2 = 0.42$** |
| NAc shell *fos*+ cells co-expressing CtB [% total *fos*] | $F_{(1,11)} = 0.73$,<br>$p = 0.41$ | **$F_{(1,11)} = 5.36$,**<br>**$p = 0.04$, $\eta_p^2 = 0.32$** | **$F_{(1,11)} = 5.03$,**<br>**$p = 0.04$, $\eta_p^2 = 0.31$** |

## DISCUSSION

Here, we demonstrated that inactivation of the VP, either through pharmacological activation of GABA_A_ receptors in the VP or through chemogenetic stimulation of NAc^GABA^ terminals in the VP, reduced the expression of social play behaviors in both male and female juvenile rats. We further showed that this behavioral effect was specific to social play because there were no changes in other social behaviors and no decreases in locomotor activity. Thus, our findings demonstrate that activation of the VP, through reduced NAc inhibitory input, is necessary for the typical expression of social play behavior in male and female juvenile rats. Additionally, we showed that the equal expression of social play behavior in males and females is associated with a female-specific increase in NAc shell activation and a male-specific decrease in activation of the NAc shell neurons projecting to the VP. These sex-specific changes in NAc activity following social play exposure eliminated baseline sex differences in NAc activity. Based on these findings, we propose a model in which sex-specific modulation of NAc inhibitory input to the VP facilitates activation of the VP that is necessary for the typical and equal expression of social play behavior in male and female juvenile rats.

### Activation of the VP is necessary for the typical expression of social play behavior in male and female juvenile rats

We showed that infusion of the GABA_A_ receptor agonist muscimol into the VP decreased the expression of social play behaviors in male and female juvenile rats. A previous study demonstrated that infusion of the GABA_A_ receptor agonist muscimol into the VP decreased the expression of maternal behaviors in rats (Numan et al., 2005). Together, these findings suggests that activation of the VP is necessary for the expression of highly motivated and rewarding social behaviors across the lifespan.

Although the VP receives inputs from regions such as the NAc and VTA (Williams et al., 1977; Groenewegen and Russchen, 1984; Zhou et al., 2022), the VP itself encodes reward values of external stimuli in adult male rats (Richard et al., 2016; Ottenheimer et al., 2018; Vento and Jhou, 2020), suggesting activity within the VP can modulate the reward incentive of stimuli and alter the expression of rewarding behaviors accordingly. Although these studies utilized adult rodents, findings from the current study suggest that modulation of VP activity may be involved in encoding reward value in juvenile rodents as well. In support, exposure to social play altered activation of *fos*+ cells in the VP in male juvenile rats (Lee et al., 2024) compared to male juvenile rats that were not exposed to social play. Future studies should investigate *in vivo* patterns of VP neuronal firing in juvenile rats as they are engaging in social play behavior to better understand the role of VP activation in regulating social play behavior.

### Inhibition of NAc^GABA^ inputs to the VP is necessary for the typical expression of social play behavior in male and female juvenile rats

We found that chemogenetic stimulation of NAc^GABA^ terminals in the VP reduced social play behaviors and VP cell activation in male and female juvenile rats. These findings suggest that reduced activation of the NAc to VP pathway is necessary for the typical expression of social play behaviors in both sexes. Our findings are in line with previous studies on rewarding behaviors showing that optogenetic stimulation of NAc terminals in the VP decreased sucrose intake in adult female rats (Chometton et al., 2020), while chemogenetic inhibition of NAc terminals in the VP increased the effort to obtain a food reward in adult male and female mice (Gallo et al., 2018). Together, these studies provide evidence that reduced activation of NAc inputs to the VP is necessary for the expression of rewarding behaviors. Our study is the first to indicate that this is true for juvenile-typical rewarding social behavior. Combined with the findings from adult rodents, it is plausible that inhibition of NAc inputs to the VP may be necessary for rewarding behaviors, social or non-social. Future studies should elucidate the involvement of this pathway in the regulation of a broad range of rewarding behaviors.

A methodological limitation of the current study is that DREADDs transduction was observed in both the NAc core and shell. Therefore, it is not possible to differentiate whether the behavioral changes observed in the current study are due to inhibition of NAc core or NAc shell inputs to the VP. It is also possible that inhibition of both inputs are required to disinhibit the VP to modulate social play behavior. The NAc core projects primarily to the dorsolateral region of the VP (Zahm and Brog, 1992; Zahm et al., 1996), while the NAc shell projects primarily to the ventromedial (Zahm and Heimer, 1990; Heimer et al., 1991) and ventrolateral (Zahm and Brog, 1992; Zahm et al., 1996) regions of the VP. Therefore, distinct experimental modulation of NAc core and shell inputs to the VP will help to better understand which of these inputs are involved in the regulation of social play behavior in juvenile rats.

### Social play exposure is associated with increased activation of the NAc core in both sexes and female-specific increased activation of the NAc shell

We found that social play exposure is associated with an increase in the number of *fos*+ cells in the NAc core of male and female juvenile rats. This is in line with a previous study reporting that social play exposure is associated with an increase in the number of Fos-immunoreactive cells in the NAc core of male juvenile rats (females were not included; (van Kerkhof et al., 2014). On the other hand, we found that only females showed an increase in the number of *fos*+ cells in the NAc shell upon social play exposure. The observed lack of a change in NAc shell activation in males exposed to social play is in contrast with a previous study showing an increase in the number of Fos-immunoreactive cells in the NAc shell of male juvenile rats exposed to social play (van Kerkhof et al., 2014).This discrepancy could be due to several methodological differences between van Kerkhof et al. (2014) and our study, including testing in the light versus dark phase, exposure to a familiar versus novel stimulus rat, and quantifying c-Fos-positive cell density versus number of fos mRNA-expressing cells. Yet, these methodological differences did not have an effect on a social play-associated increase in activation of the NAc core, suggesting a highly robust involvement of the NAc core in social play behavior in juvenile rats.

We hypothesized that activation of the VP is necessary for the expression of social play through inhibition of the NAc to VP pathway. Therefore, we would have expected to find a decrease, rather than an increase, in NAc cell activation in response to social play. However, the NAc projects to brain regions other than the VP as well as to local circuits within the NAc (Usuda et al., 1998). Accordingly, it is unclear whether an increase in the number of *fos*+ cells reflects an increase in local excitatory or inhibitory transmissions in the NAc. This is an important distinction because it could be that NAc GABAergic interneurons were activated in response to social play exposure, which suppress activation of NAc MSNs (Qi et al., 2016; Wright et al., 2017; Yu et al., 2017). This suggestion would be in line with the observation that pharmacological inactivation of the NAc core increased social play behaviors in male juvenile rats (females were not tested; (van Kerkhof et al., 2013). The same study also showed that pharmacological inactivation of the NAc shell did not alter social play behaviors in males. Our finding of a female-specific increase in activation of the NAc shell suggest that this mechanism could be different in females. Future experiments could determine the phenotype (interneurons versus output neurons) of activated cells in the NAc core and shell to reveal how NAc activity is involved in social play behavior in males and females.

### Social play exposure is associated with male-specific reduction in activation of NAc core and shell neurons projecting to the VP

We found that exposure to social play was associated with reduced activation of NAc core and shell neurons projecting to the VP in male juvenile rats. Moreover, we previously reported that social play exposure was associated with an increase in VP neuronal activation in male juvenile rats (Lee et al., 2024). Together, these findings support our hypothesis, at least in males, that activation of the VP is necessary for the expression of social play through inhibition of the NAc to VP pathway. However, females showed a very different activation pattern of the NAc to VP pathway. There was a strong trend (p = 0.053) toward increased activation of NAc core inputs to the VP of females exposed to social play without a change in activation of NAc shell neurons projecting to the VP. Subsequently, we previously observed that social play exposure was associated with no change in VP activation in female juvenile rats (Lee et al., 2024). It should be noted that different cohorts of rats in the “No Social Play” and “Social Play” groups were used to analyze the NAc core and shell separately as the NAc core and shell were analyzed on different planes. Therefore, there are noted inconsistencies in the number of subjects included in the analysis of the NAc core and shell. Regardless, these findings, combined with the observed female-specific increase in NAc shell activation in response to social play exposure, suggest sex differences in the roles of the NAc core and shell in regulating social play behavior. As mentioned earlier, the NAc core and shell innervate different regions of the VP and these subregions of the VP also have distinct outputs (Carlsen et al., 1985; Haber et al., 1985; Kalivas et al., 1993; Mascagni and McDonald, 2009). Based on the observed sex-specific changes in activation of VP-projecting cells in the NAc core and shell in combination with findings of chemogenetic inhibition of the NAc to VP pathway, it is possible that juvenile males and females may engage different subregional pathways between the NAc and VP in order exhibit similar levels of social play. Future studies could test this hypothesis.

We showed that females had lower baseline activation of NAc core and shell neurons projecting to the VP compared to males. One may expect that this would result in higher baseline VP neuronal activity in females versus males. However, males and females showed similar baseline VP neuronal activation (Lee et al., 2024). Thus, a sex difference in activation of NAc inputs to the VP may support similar VP neuronal activation under baseline conditions in juvenile rats. Although we are not aware of other studies reporting sex differences in NAc inputs to the VP, there is evidence suggesting sex differences in afferents to the NAc, thereby potentially modulating NAc inputs to the VP in a sex-specific manner. For example, adult female rats showed higher spine density in the NAc core and a trend towards higher spine density in the NAc shell compared to males (Forlano and Woolley, 2010; Wissman et al., 2012). Adult female rats also showed larger spine heads in the NAc core and shell compared to males (Forlano and Woolley, 2010). These findings suggest that females receive denser inputs from upstream regions than males, which may result in sex differences in baseline NAc activity, including those to downstream targets. Whether a similar mechanism exists at the juvenile age remains to be determined.

### Sex-specific activation of the NAc and NAc to VP pathway underlie similar levels of social play behaviors in male and female juvenile rats

In our study, male and female juvenile Wistar and Long Evans rats showed similar levels of social play behaviors. This is in line with previous studies from our lab and others using a paradigm in which rats are isolated for 24 hours and then tested for social play behavior in their home cage (Veenema et al., 2013; Paul et al., 2014; Bredewold et al., 2015, 2018; Northcutt and Nwankwo, 2018; Reppucci et al., 2018, 2020; Lee et al., 2024). Yet, these similar social play levels are associated with sex-specific activation of the NAc and NAc to VP pathway upon social play exposure. This too corresponds with previous work from our lab in which equal levels of social play behaviors are associated with sex-specific changes or function of various neurotransmitter systems (including vasopressin, oxytocin, dopamine, glutamate) across brain regions (Veenema et al., 2013; Bredewold et al., 2014, 2015, 2018; Lee et al., 2024). Several other studies show that sex-specific involvement of the NAc supports the similar expression of other rewarding social and non-social behaviors. For example, cocaine exposure was associated with a higher release of dopamine in the NAc in adult male rats compared to adult female rats (Cummings et al., 2014). Similarly, adult male mice displayed higher extracellular concentration of dopamine in the NAc shell under baseline and following social defeat stress compared to adult female mice (Campi et al., 2014). Additionally, infusions of the D1R agonist SKF38393 into the NAc shell reduced social interaction time in adult female mice while no change was observed in adult male mice (Campi et al., 2014). These studies along with the current study show that in juvenile and adult rodents, the NAc is engaged differently in males versus females and this mechanism may be crucial to support the similar expression of social and non-social rewarding behaviors.

## Conclusions

We show that activation of the VP is necessary for the typical expression of social play behavior in both male and female juvenile rats, suggesting that disinhibition of the VP may be an important mechanism to support social play behavior. Indeed, we show that inhibition of the NAc^GABA^ to VP pathway, thereby disinhibiting the VP, is necessary for the expression of typical social play levels in both sexes. Yet, we find that the underlying mechanisms are sex-specific resulting in the elimination of a baseline sex difference in the activation of the NAc and the NAc to VP pathway. Together, these findings provide the first evidence of the functional involvement of the NAc to VP pathway in regulating social play behavior in juvenile rats. It is likely that the NAc and VP are part of a larger reward circuitry regulating social play behavior. Future investigations could elucidate how this reward circuitry regulates the equal expression of social play behavior through sex-specific mechanisms in juvenile rats.

## Acknowledgements

We would like to thank members of the Veenema lab for critically reading an earlier version of this manuscript and animal caretakers for excellent animal care.

## Author Contributions

JDAL performed experiments, developed the concept, analyzed the data, wrote the manuscript, and obtained funding. DNA and ICO performed experiments. SMB performed experiments and provided input to the manuscript. AHV developed the concept, supervised the project, wrote the manuscript, and obtained funding.

## References

Achterberg EJM, van Kerkhof LWM, Servadio M, van Swieten MMH, Houwing DJ, Aalderink M, Driel NV, Trezza V, Vanderschuren LJMJ (2016a) Contrasting roles of dopamine and noradrenaline in the motivational properties of social play behavior in rats. Neuropsychopharmacology 41:858–868.

Achterberg EJM, van Swieten MMH, Driel NV, Trezza V, Vanderschuren LJMJ (2016b) Dissociating the role of endocannabinoids in the pleasurable and motivational properties of social play behaviour in rats. Pharmacol Res 110:151–158.

Achterberg EJM, van Swieten MMH, Houwing DJ, Trezza V, Vanderschuren LJMJ (2019) Opioid modulation of social play reward in juvenile rats. Neuropharmacology 159:107332.

Bekoff M (1974) Social play in coyotes, wolves, and dogs. Bioscience 24:225–230.

Berridge KC, Kringelbach ML (2015) Pleasure systems in the brain. Neuron 86:646–664.

Bredewold R, Nascimento NF, Ro GS, Cieslewski SE, Reppucci CJ, Veenema AH (2018) Involvement of dopamine, but not norepinephrine, in the sex-specific regulation of juvenile socially rewarding behavior by vasopressin. Neuropsychopharmacology 43:2109–2117.

Bredewold R, Schiavo JK, van der Hart M, Verreij M, Veenema AH (2015) Dynamic changes in extracellular release of GABA and glutamate in the lateral septum during social play behavior in juvenile rats: Implications for sex-specific regulation of social play behavior. Neuroscience 307:117–127.

Bredewold R, Smith CJW, Dumais KM, Veenema AH (2014) Sex-specific modulation of juvenile social play behavior by vasopressin and oxytocin depends on social context. Front Behav Neurosci 8:216.

Brog JS, Salyapongse A, Deutch AY, Zahm DS (1993) The patterns of afferent innervation of the core and shell in the “accumbens” part of the rat ventral striatum: immunohistochemical detection of retrogradely transported fluoro-gold. J Comp Neurol 338:255–278.

Buggey T, Hoomes G, Sherberger ME, Williams S (2011) Facilitating Social Initiations of Preschoolers With Autism Spectrum Disorders Using Video Self-Modeling. Focus Autism Other Dev Disabl 26:25–36.

Calcagnetti DJ, Schechter MD (1992) Place conditioning reveals the rewarding aspect of social interaction in juvenile rats. Physiol Behav 51:667–672.

Campi KL, Greenberg GD, Kapoor A, Ziegler TE, Trainor BC (2014) Sex differences in effects of dopamine D1 receptors on social withdrawal. Neuropharmacology 77:208–216.

Carlezon WA, Thomas MJ (2009) Biological substrates of reward and aversion: a nucleus accumbens activity hypothesis. Neuropharmacology 56 Suppl 1:122–132.

Carlsen J, Záborszky L, Heimer L (1985) Cholinergic projections from the basal forebrain to the basolateral amygdaloid complex: a combined retrograde fluorescent and immunohistochemical study. J Comp Neurol 234:155–167.

Chometton S, Guèvremont G, Seigneur J, Timofeeva E, Timofeev I (2020) Projections from the nucleus accumbens shell to the ventral pallidum are involved in the control of sucrose intake in adult female rats. Brain Struct Funct 225:2815–2839.

Cummings JA, Jagannathan L, Jackson LR, Becker JB (2014) Sex differences in the effects of estradiol in the nucleus accumbens and striatum on the response to cocaine: neurochemistry and behavior. Drug Alcohol Depend 135:22–28.

Farrell MR, Esteban JSD, Faget L, Floresco SB, Hnasko TS, Mahler SV (2021) Ventral pallidum GABA neurons mediate motivation underlying risky choice. J Neurosci 41:4500–4513.

Farrell MR, Ye Q, Xie Y, Esteban JSD, Mahler SV (2022) Ventral pallidum GABA neurons bidirectionally control opioid relapse across rat behavioral models. Addiction Neuroscience 3:100026.

Forlano PM, Woolley CS (2010) Quantitative analysis of pre- and postsynaptic sex differences in the nucleus accumbens. J Comp Neurol 518:1330–1348.

Gallo EF, Meszaros J, Sherman JD, Chohan MO, Teboul E, Choi CS, Moore H, Javitch JA, Kellendonk C (2018) Accumbens dopamine D2 receptors increase motivation by decreasing inhibitory transmission to the ventral pallidum. Nat Commun 9:1086.

Gibson GD, Prasad AA, Jean-Richard-Dit-Bressel P, Yau JOY, Millan EZ, Liu Y, Campbell EJ, Lim J, Marchant NJ, Power JM, Killcross S, Lawrence AJ, McNally GP (2018) Distinct Accumbens Shell Output Pathways Promote versus Prevent Relapse to Alcohol Seeking. Neuron 98:512–520.e6.

Groenewegen HJ, Russchen FT (1984) Organization of the efferent projections of the nucleus accumbens to pallidal, hypothalamic, and mesencephalic structures: a tracing and immunohistochemical study in the cat. J Comp Neurol 223:347–367.

Haber SN, Groenewegen HJ, Grove EA, Nauta WJ (1985) Efferent connections of the ventral pallidum: evidence of a dual striato pallidofugal pathway. J Comp Neurol 235:322–335.

Heimer L, Zahm DS, Churchill L, Kalivas PW, Wohltmann C (1991) Specificity in the projection patterns of accumbal core and shell in the rat. Neuroscience 41:89–125.

Holmes E, Willoughby T (2005) Play behaviour of children with autism spectrum disorders. J Intellect Dev Dis 30:156–164.

Ikemoto S, Panksepp J (1992) The effects of early social isolation on the motivation for social play in juvenile rats. Dev Psychobiol 25:261–274.

Jones DL, Mogenson GJ (1980) Nucleus accumbens to globus pallidus GABA projection: electrophysiological and iontophoretic investigations. Brain Res 188:93–105.

Kalivas PW, Churchill L, Klitenick MA (1993) GABA and enkephalin projection from the nucleus accumbens and ventral pallidum to the ventral tegmental area. Neuroscience 57:1047–1060.

Lee JDA, Reppucci CJ, Huez EDM, Bredewold R, Veenema AH (2024) Sex differences in the structure and function of the vasopressin system in the ventral pallidum are associated with the sex-specific regulation of social play behavior in juvenile rats. Horm Behav 163:105563.

Marquardt AE, VanRyzin JW, Fuquen RW, McCarthy MM (2022) Social play experience in juvenile rats is indispensable for appropriate socio-sexual behavior in adulthood in males but not females. Front Behav Neurosci 16:1076765.

Mascagni F, McDonald AJ (2009) Parvalbumin-immunoreactive neurons and GABAergic neurons of the basal forebrain project to the rat basolateral amygdala. Neuroscience 160:805–812.

Morgan JI, Curran T (1991) Stimulus-transcription coupling in the nervous system: involvement of the inducible proto-oncogenes fos and jun. Annu Rev Neurosci 14:421–451.

Northcutt KV, Nwankwo VC (2018) Sex differences in juvenile play behavior differ among rat strains. Dev Psychobiol 60:903–912.

Numan M, Numan MJ, Pliakou N, Stolzenberg DS, Mullins OJ, Murphy JM, Smith CD (2005) The effects of D1 or D2 dopamine receptor antagonism in the medial preoptic area, ventral pallidum, or nucleus accumbens on the maternal retrieval response and other aspects of maternal behavior in rats. Behav Neurosci 119:1588–1604.

O’Donnell P, Grace AA (1993) Physiological and morphological properties of accumbens core and shell neurons recorded in vitro. Synapse 13:135–160.

Ottenheimer D, Richard JM, Janak PH (2018) Ventral pallidum encodes relative reward value earlier and more robustly than nucleus accumbens. Nat Commun 9:4350.

Panksepp J (1981) The ontogeny of play in rats. Dev Psychobiol 14:327–332.

Paul MJ, Terranova JI, Probst CK, Murray EK, Ismail NI, de Vries GJ (2014) Sexually dimorphic role for vasopressin in the development of social play. Front Behav Neurosci 8:58.

Paxinos, G. and Watson, C. (2007) The Rat Brain in Stereotaxic Coordinates. 6th Edition, Academic Press, San Diego.

Pellis SM, Pellis VC, Burke CJ, Stark RA, Ham JR, Euston DR, Achterberg EJM (2022) Measuring play fighting in rats: A multilayered approach. Curr Protoc 2:e337.

Pellis SM, Pellis VC (1987) Play-fighting differs from serious fighting in both target of attack and tactics of fighting in the laboratory ratRattus norvegicus. Aggress Behav 13:227–242.

Pellis SM, Pellis VC (1990) Differential rates of attack, defense, and counterattack during the developmental decrease in play fighting by male and female rats. Dev Psychobiol 23:215–231.

Qi J, Zhang S, Wang H-L, Barker DJ, Miranda-Barrientos J, Morales M (2016) VTA glutamatergic inputs to nucleus accumbens drive aversion by acting on GABAergic interneurons. Nat Neurosci 19:725–733.

Reppucci CJ, Gergely CK, Bredewold R, Veenema AH (2020) Involvement of orexin/hypocretin in the expression of social play behaviour in juvenile rats. International Journal of Play:1–20.

Reppucci CJ, Gergely CK, Veenema AH (2018) Activation patterns of vasopressinergic and oxytocinergic brain regions following social play exposure in juvenile male and female rats. J Neuroendocrinol.

Richard JM, Ambroggi F, Janak PH, Fields HL (2016) Ventral Pallidum Neurons Encode Incentive Value and Promote Cue-Elicited Instrumental Actions. Neuron 90:1165–1173.

Root DH, Melendez RI, Zaborszky L, Napier TC (2015) The ventral pallidum: Subregion-specific functional anatomy and roles in motivated behaviors. Prog Neurobiol 130:29–70.

Salamone JD, Correa M (2002) Motivational views of reinforcement: implications for understanding the behavioral functions of nucleus accumbens dopamine. Behav Brain Res 137:3–25.

Scott E, Panksepp J (2003) Rough-and-tumble play in human children. Aggress Behav 29:539–551.

Sharpe MJ, Marchant NJ, Whitaker LR, Richie CT, Zhang YJ, Campbell EJ, Koivula PP, Necarsulmer JC, Mejias-Aponte C, Morales M, Pickel J, Smith JC, Niv Y, Shaham Y, Harvey BK, Schoenbaum G (2017) Lateral Hypothalamic GABAergic Neurons Encode Reward Predictions that Are Relayed to the Ventral Tegmental Area to Regulate Learning. Curr Biol 27:2089–2100.e5.

Smith KS, Berridge KC, Aldridge JW (2011) Disentangling pleasure from incentive salience and learning signals in brain reward circuitry. Proc Natl Acad Sci USA 108:E255–64.

Supekar K, Kochalka J, Schaer M, Wakeman H, Qin S, Padmanabhan A, Menon V (2018) Deficits in mesolimbic reward pathway underlie social interaction impairments in children with autism. Brain 141:2795–2805.

Swerdlow NR, Braff DL, Geyer MA (1990) GABAergic projection from nucleus accumbens to ventral pallidum mediates dopamine-induced sensorimotor gating deficits of acoustic startle in rats. Brain Res 532:146–150.

Thor DH, Holloway WR (1984a) Developmental analyses of social play behavior in juvenile rats. Bull Psychon Soc 22:587–590.

Thor DH, Holloway WR (1984b) Social play in juvenile rats: a decade of methodological and experimental research. Neurosci Biobehav Rev 8:455–464.

Trezza V, Damsteegt R, Vanderschuren LJMJ (2009) Conditioned place preference induced by social play behavior: parametrics, extinction, reinstatement and disruption by methylphenidate. Eur Neuropsychopharmacol 19:659–669.

Usuda I, Tanaka K, Chiba T (1998) Efferent projections of the nucleus accumbens in the rat with special reference to subdivision of the nucleus: biotinylated dextran amine study. Brain Res 797:73–93.

van den Berg CL, Hol T, Van Ree JM, Spruijt BM, Everts H, Koolhaas JM (1999) Play is indispensable for an adequate development of coping with social challenges in the rat. Dev Psychobiol 34:129–138.

van Kerkhof LWM, Damsteegt R, Trezza V, Voorn P, Vanderschuren LJMJ (2013) Social play behavior in adolescent rats is mediated by functional activity in medial prefrontal cortex and striatum. Neuropsychopharmacology 38:1899–1909.

van Kerkhof LWM, Trezza V, Mulder T, Gao P, Voorn P, Vanderschuren LJMJ (2014) Cellular activation in limbic brain systems during social play behaviour in rats. Brain Struct Funct 219:1181–1211.

Veenema AH, Bredewold R, De Vries GJ (2013) Sex-specific modulation of juvenile social play by vasopressin. Psychoneuroendocrinology 38:2554–2561.

Veenema AH, Neumann ID (2009) Maternal separation enhances offensive play-fighting, basal corticosterone and hypothalamic vasopressin mRNA expression in juvenile male rats. Psychoneuroendocrinology 34:463–467.

Vento PJ, Jhou TC (2020) Bidirectional valence encoding in the ventral pallidum. Neuron 105:766–768.

Walaas I, Fonnum F (1979) The distribution and origin of glutamate decarboxylase and choline acetyltransferase in ventral pallidum and other basal forebrain regions. Brain Res 177:325–336.

Williams DJ, Crossman AR, Slater P (1977) The efferent projections of the nucleus accumbens in the rat. Brain Res 130:217–227.

Wissman AM, May RM, Woolley CS (2012) Ultrastructural analysis of sex differences in nucleus accumbens synaptic connectivity. Brain Struct Funct 217:181–190.

Wright WJ, Schlüter OM, Dong Y (2017) A Feedforward Inhibitory Circuit Mediated by CB1-Expressing Fast-Spiking Interneurons in the Nucleus Accumbens. Neuropsychopharmacology 42:1146–1156.

Yu J, Yan Y, Li K-L, Wang Y, Huang YH, Urban NN, Nestler EJ, Schlüter OM, Dong Y (2017) Nucleus accumbens feedforward inhibition circuit promotes cocaine self-administration. Proc Natl Acad Sci USA 114:E8750–E8759.

Zahm DS, Brog JS (1992) On the significance of subterritories in the “accumbens” part of the rat ventral striatum. Neuroscience 50:751–767.

Zahm DS, Heimer L (1990) Two transpallidal pathways originating in the rat nucleus accumbens. J Comp Neurol 302:437–446.

Zahm DS, Williams E, Wohltmann C (1996) Ventral striatopallidothalamic projection: IV. Relative involvements of neurochemically distinct subterritories in the ventral pallidum and adjacent parts of the rostroventral forebrain. J Comp Neurol 364:340–362.

Zhou W-L, Kim K, Ali F, Pittenger ST, Calarco CA, Mineur YS, Ramakrishnan C, Deisseroth K, Kwan AC, Picciotto MR (2022) Activity of a direct VTA to ventral pallidum GABA pathway encodes unconditioned reward value and sustains motivation for reward. Sci Adv 8:eabm5217.

