## Supplemental Materials for "The nucleus accumbens to ventral pallidum pathway regulates social play behavior via sex-specific mechanisms in juvenile rats"

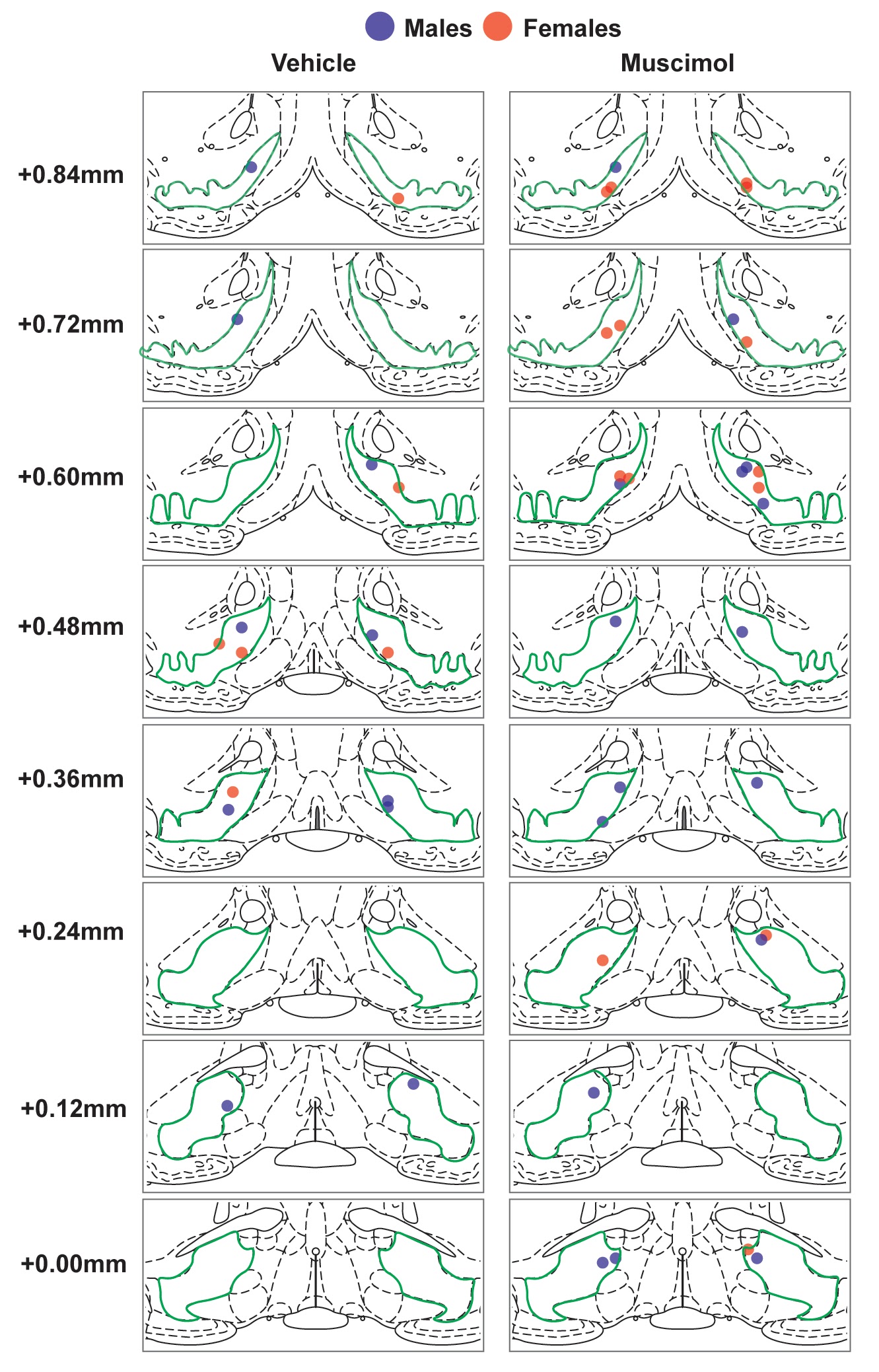

**Supplementary Figure 1.** **Experiment 1: Bilateral cannula placements in the VP** (based on the central placement of charcoal that was injected as marker) for vehicle and muscimol microinfusions shown on modified rat brain atlas templates (Paxinos and Watson, 2007). Ventral pallidum is outlined in green; numbers to the left of the atlas templates represent distance from Bregma. Ventral pallidum is outlined in green; numbers to the left of the atlas templates represent distance from Bregma.

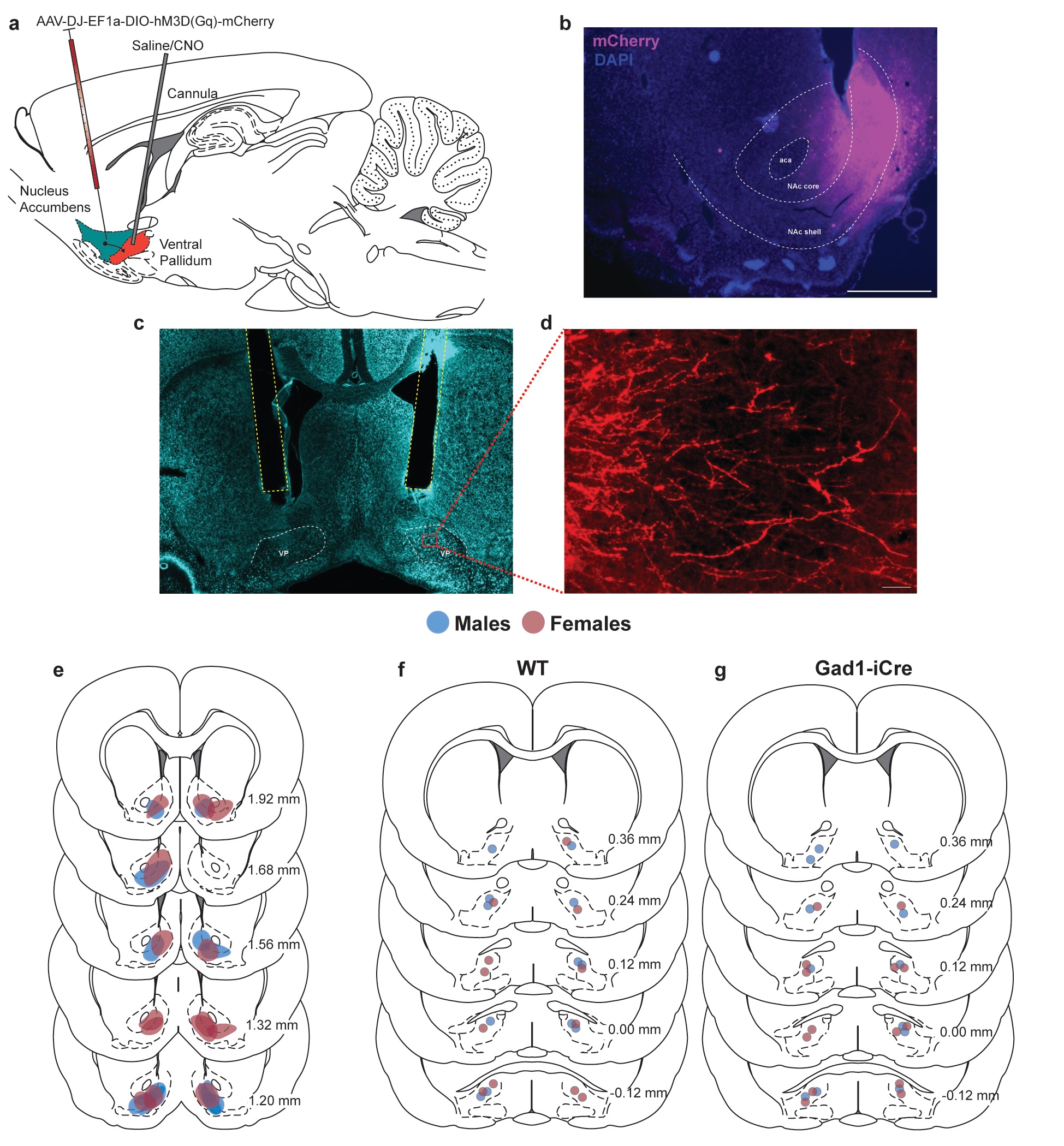

**Supplementary Figure 2. Experiment 2: DREADDs expression in the NAc and VP and cannulation of the VP in juvenile male and female Gad1-iCre rats.** (a) Schematic illustrating bilateral infusions of the cre-dependent excitatory DREADDs (AAV-DJ-EF1a-DIO-hM3D(Gq)-mCherry) targeting the NAc and bilateral cannula implantation targeting the VP. (b) DREADDs expression (purple) is localized within the borders of the NAc (NAc subregions outlined in white). Scale bar = 500 μm. (c) Fluorescent Nissl photomicrograph of a coronal brain section depicting bilateral cannula placement (outlined in yellow) targeting the VP. Cannulas are implanted 2 mm above the VP (outlined in white). (d) Representative photomicrograph of DREADDs-positive fibers (shown in red) in the VP originating from NAc^GABA^ cells. Scale bar = 15 μm. (e) Coronal sections depicting the center of DREADDs infusions in Gad1-iCre males and females. Coronal sections depicting bilateral cannula placements for WT (f) and Gad1-iCre (g) males and females.Numbers at the right refer to distance from Bregma (in mm; Paxinos and Watson, 2007).

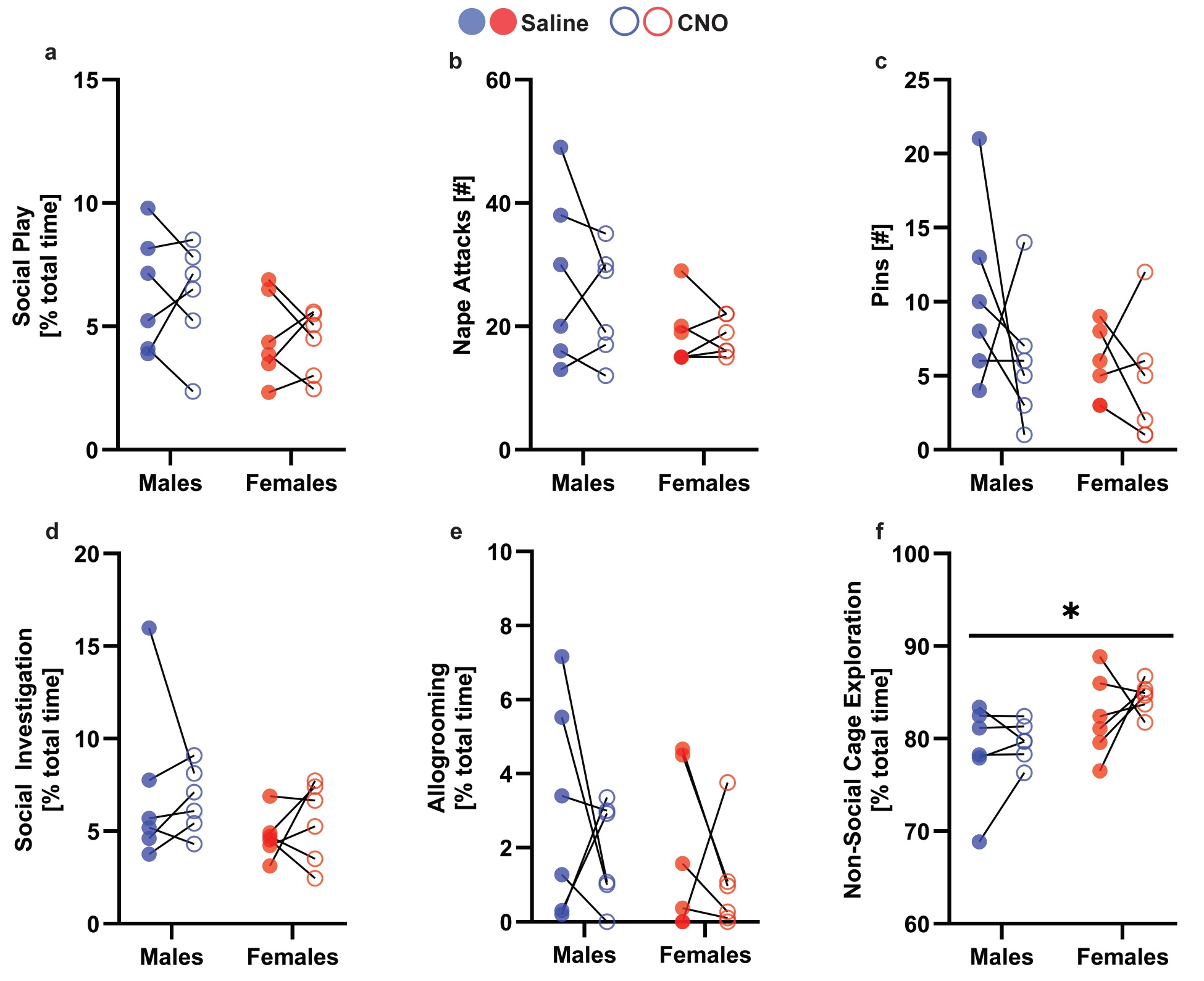

**Supplementary Figure 3. Experiment 2A: CNO by itself did not alter any behaviors during the social play test in juvenile male and female wildtype rats.** Males and females showed a similar duration of social play (a), social investigation (d), and allogrooming (e) when treated with saline and CNO. In addition, wildtype rats showed a similar number of nape attacks (b) and pins (b) following saline and CNO treatments. However, females showed a higher duration of non-social cage exploration compared to males, regardless of drug treatment (f). *p < 0.05, main effect repeated measures ANOVA.

**Supplementary Table 1.** **Experiment 2A: Two-way ANOVA statistics for social and non-social behaviors of juvenile male and female wildtype rats**. Rats were given CNO or vehicle in counterbalanced order. Significant effects are indicated in **bold**.

|  | Sex | Drug Treatment | Sex x Drug |
| --- | --- | --- | --- |
| Social play duration | *F*_(1,10)_ = 3.55,  *p* = 0.08 | *F*_(1,10)_ = 0.09,  *p* = 0.76 | *F*_(1,10)_ = 0.005,  *p* = 0.94 |
| Nape attacks [#] | *F*_(1,10)_ = 2.41,  *p* = 0.15 | *F*_(1,10)_ = 0.92,  *p* = 0.35 | *F*_(1,10)_ = 0.56,  *p* = 0.47 |
| Pins [#] | *F*_(1,10)_ = 4.79,  *p* = 0.06 | *F*_(1,10)_ = 1.58,  *p* = 0.23 | *F*_(1,10)_ = 0.52,  *p* = 0.48 |
| Supine poses [#] | *F*_(1,10)_ = 0.29,  *p* = 0.59 | *F*_(1,10)_ = 4.00,  *p* = 0.07 | *F*_(1,10)_ = 1.00,  *p* = 0.34 |
| Social investigation duration | *F*_(1,10)_ = 2.01,  *p* = 0.18 | *F*_(1,10)_ = 0.02,  *p* = 0.87 | *F*_(1,10)_ = 0.45,  *p* = 0.51 |
| Allogrooming duration | *F*_(1,10)_ = 1.83,  *p* = 0.21 | *F*_(1,10)_ = 1.02,  *p* = 0.33 | *F*_(1,10)_ = 0.01,  *p* = 0.89 |
| Non-social cage exploration duration | ***F*_(1,10)_ = 7.18,**  ***p* = 0.02, ƞ_p_^2^ = 0.39** | *F*_(1,10)_ = 1.17,  *p* = 0.30 | *F*_(1,10)_ = 0.16,  *p* = 0.69 |

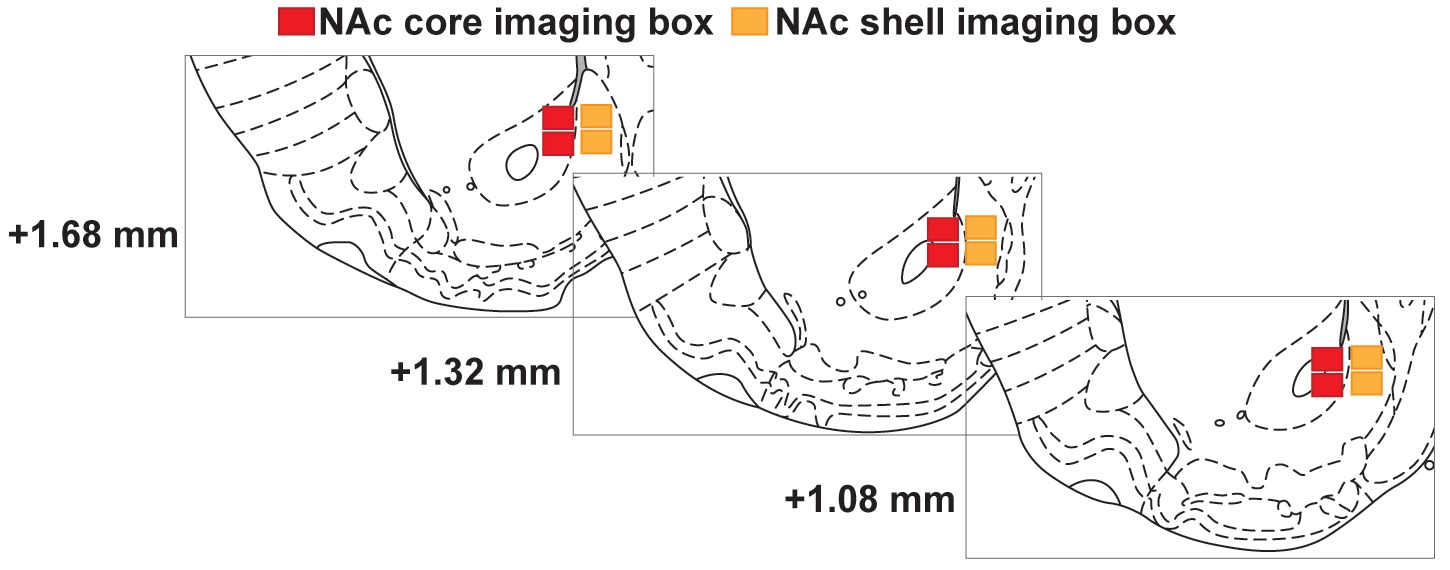

**Supplementary Figure 4.** **Experiment 3: Image sampling locations in the NAc core and NAc shell** shown on modified rat brain atlas templates (Paxinos and Watson, 2007). Numbers to the left of the atlas templates represent distance from Bregma; red boxes represent sampling locations of images taken in the NAc core; yellow boxes represent sampling locations of images taken in the NAc shell.

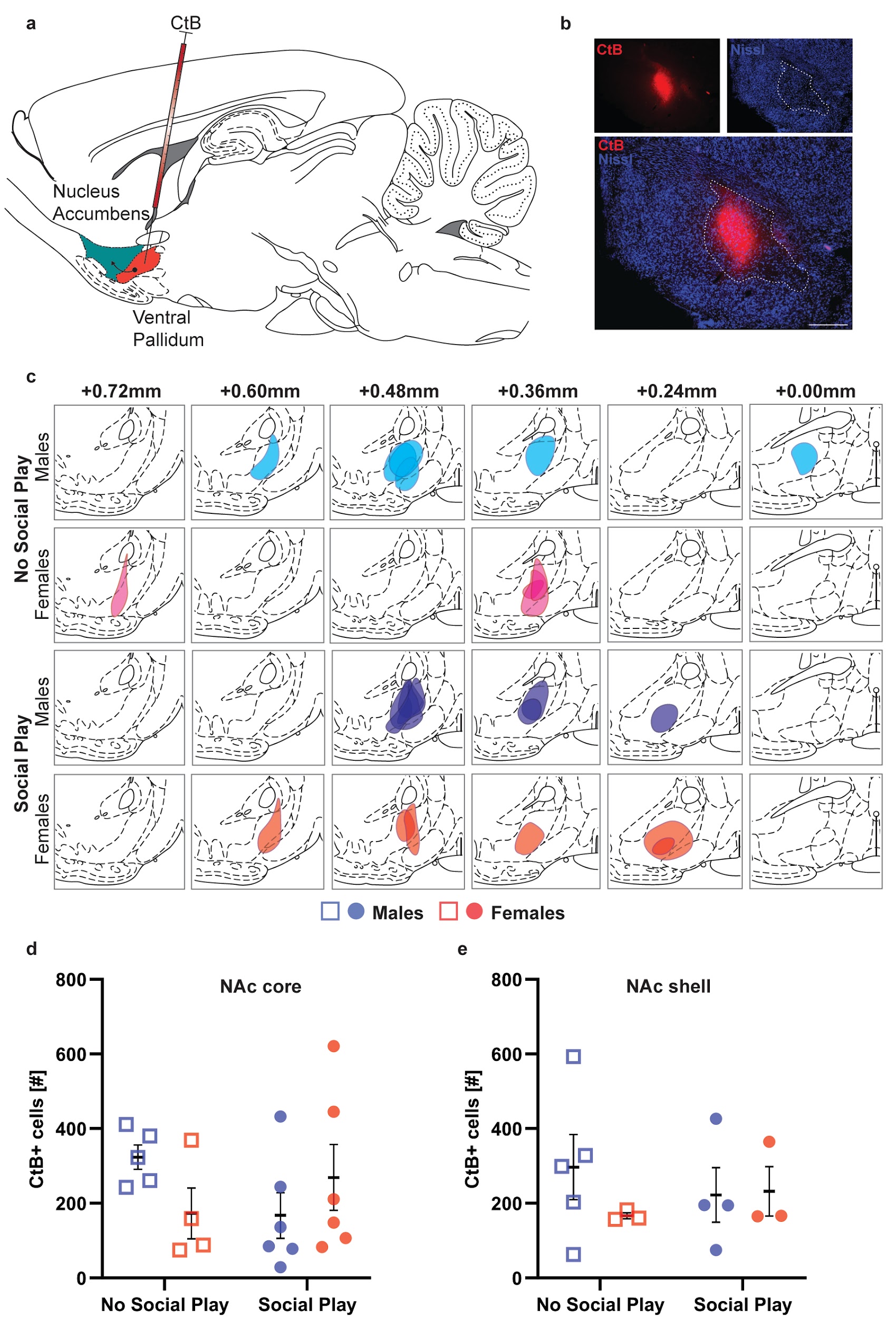

**Supplementary Figure 5. Experiment 3: No sex difference in the number of VP-projecting cells in the NAc core and shell.** (A) Schematic illustration of unilateral infusion of the retrograde tracer CtB (AF 594) targeting the VP. (B) CtB expression (red) is localized within the borders of the VP (outlined in white) defined using a fluorescent Nissl (blue). (C) Coronal sections depicting the center of CtB infusions in males and females in the “No Social Play” and “Social Play” groups. Numbers at the top refer to distance from Bregma (in mm; Paxinos and Watson, 2007). Males and females in the “No Social Play” and “Social Play” groups show similar numbers of VP-projecting cells in the NAc core (D) and NAc shell (E). Black bars indicate mean ± SEM; Scale bar = 150 μm.

**Supplementary Table 2.** **Experiment 3: Two-way ANOVA statistics for the number of CTB+ cells in the NAc core and shell** showing no significant differences between males and females nor between the “No Social Play” and “Social Play” groups.

|  | Sex | Social Play Condition | Sex x Social Play |
| --- | --- | --- | --- |
| NAc core CtB+ cells [#] | *F*_(1,17)_ = 0.0003,  p = 0.98 | *F*_(1,17)_ = 0.003,  p = 0.95 | *F*_(1,17)_ = 1.93,  p = 0.18 |
| NAc shell CtB+ cells [#] | *F*_(1,11)_ = 0.59,  p = 0.45 | *F*_(1,11)_ = 0.003,  p = 0.95 | *F*_(1,11)_ = 0.79,  p = 0.39 |

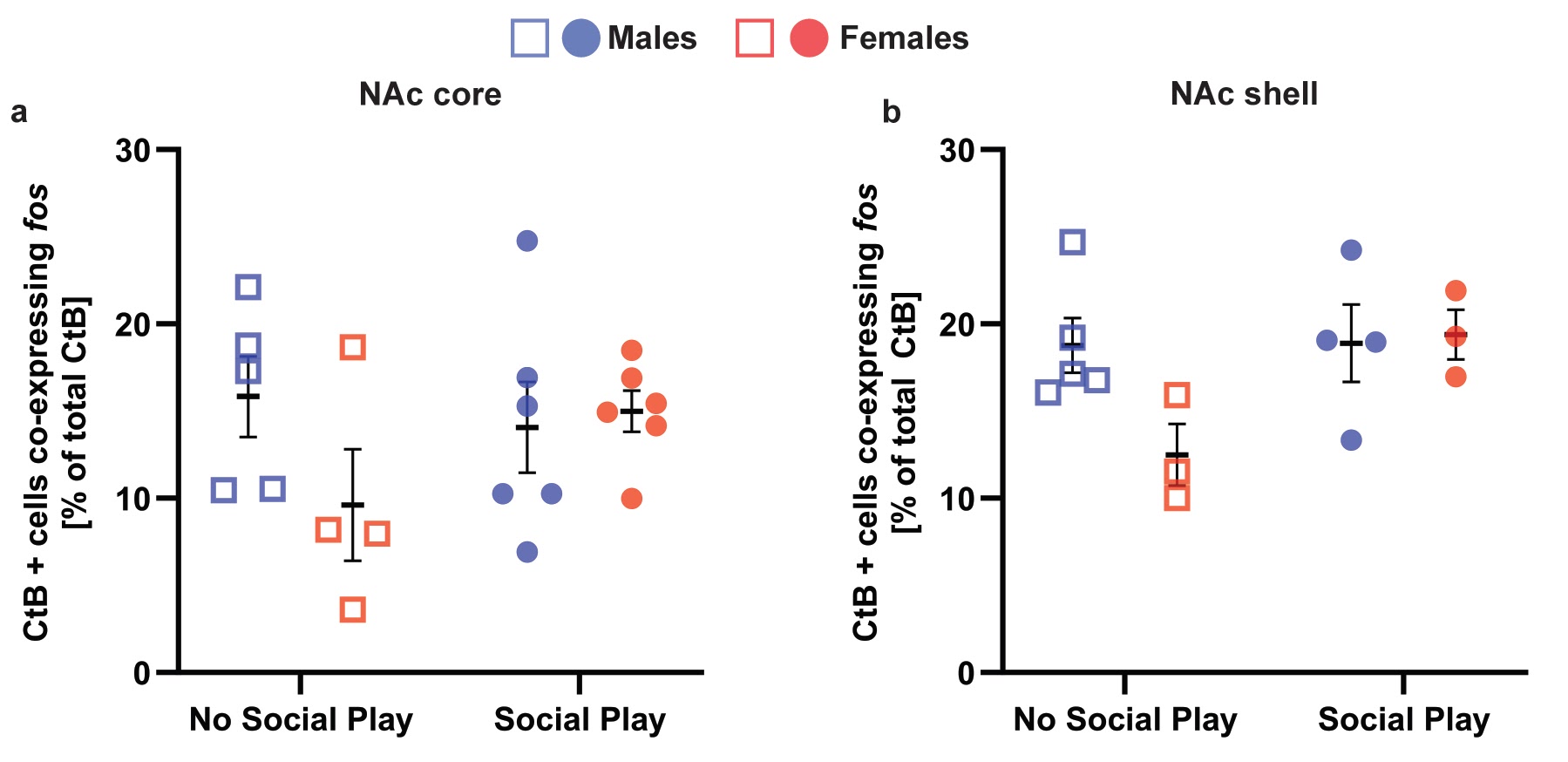

**Supplementary Figure 6. Social play exposure did not alter the proportion of recruited VP-projecting cells in the NAc core and shell in either sex.** Males and females in the “No Social Play” and “Social Play” groups showed a similar proportion of CTB+ cells that co-expressed *fos* in the NAc core (a) and NAc shell (b). Black bars indicate mean ± SEM.

**Supplementary Table 3.** **Experiment 3: Two-way ANOVA statistics for the percentage of CtB+ cells co-expressing *fos* in the NAc core and shell** showing no significant differences between males and females nor between the “No Social Play” and “Social Play” groups.

|  | Sex | Social Play Condition | Sex x Social Play |
| --- | --- | --- | --- |
| NAc core CtB+ cells co-expressing fos [% total CtB] | *F*_(1,17)_ = 1.28,  p = 0.27 | *F*_(1,17)_ = 0.59,  p = 0.45 | *F*_(1,17)_ = 2.32,  p = 0.14 |
| NAc shell CtB+ cells co-expressing fos [% total CtB] | *F*_(1,11)_ = 2.35,  p = 0.15 | *F*_(1,11)_ = 3.47,  p = 0.08 | *F*_(1,11)_ = 3.23,  p = 0.09 |

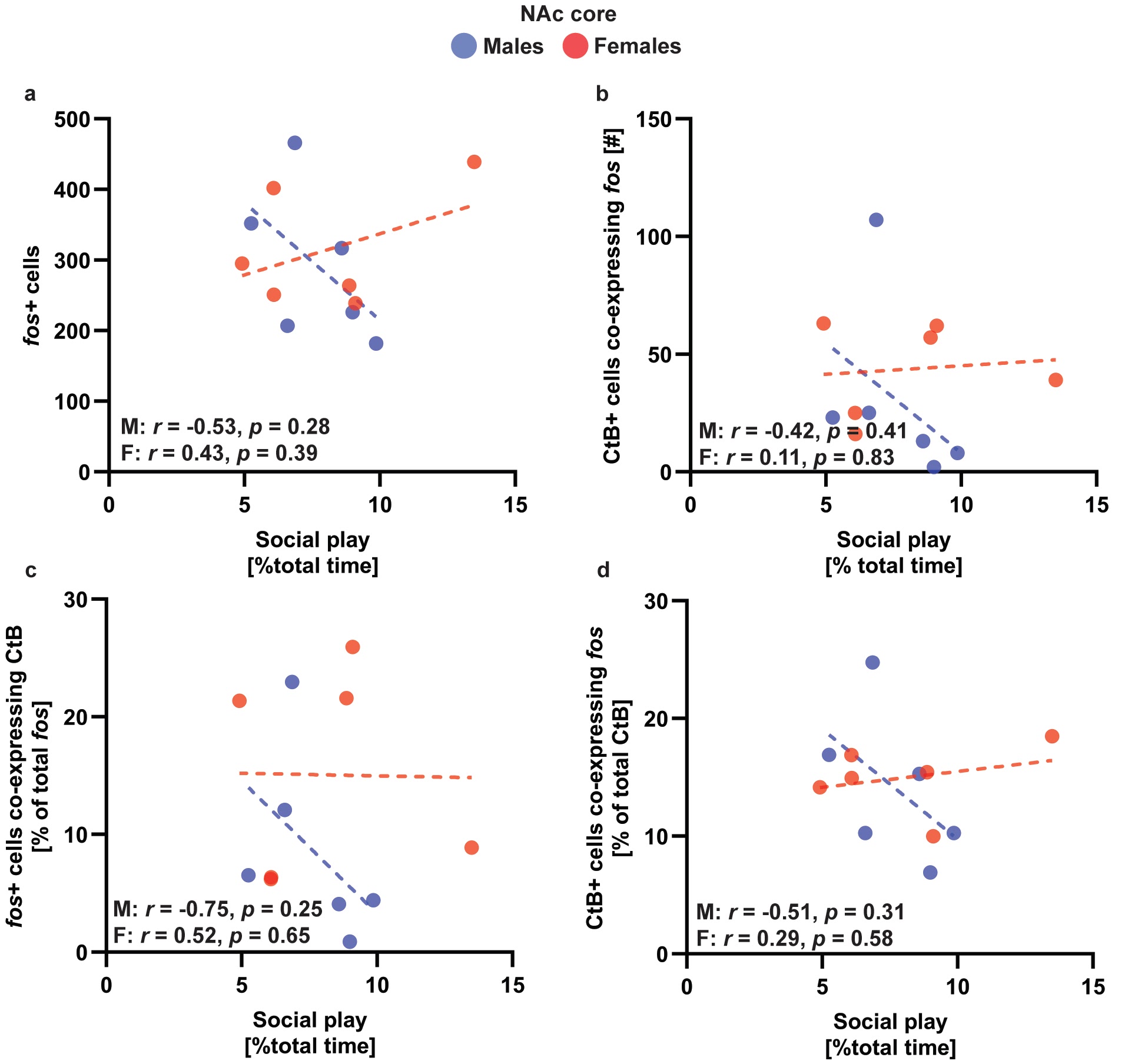

**Supplementary Figure 7. Experiment 3: Social play duration did not significantly correlate with any of the analyzed parameters in the NAc core.** The percent of time spent engaging in social play did not correlate with the number of *fos*+ cells (a), the number of CtB+ cells co-expressing *fos* (b), the proportion of recruited cells that project to the VP (c), or the proportion of VP-projecting cells activated (d) in the NAc core of either sex.

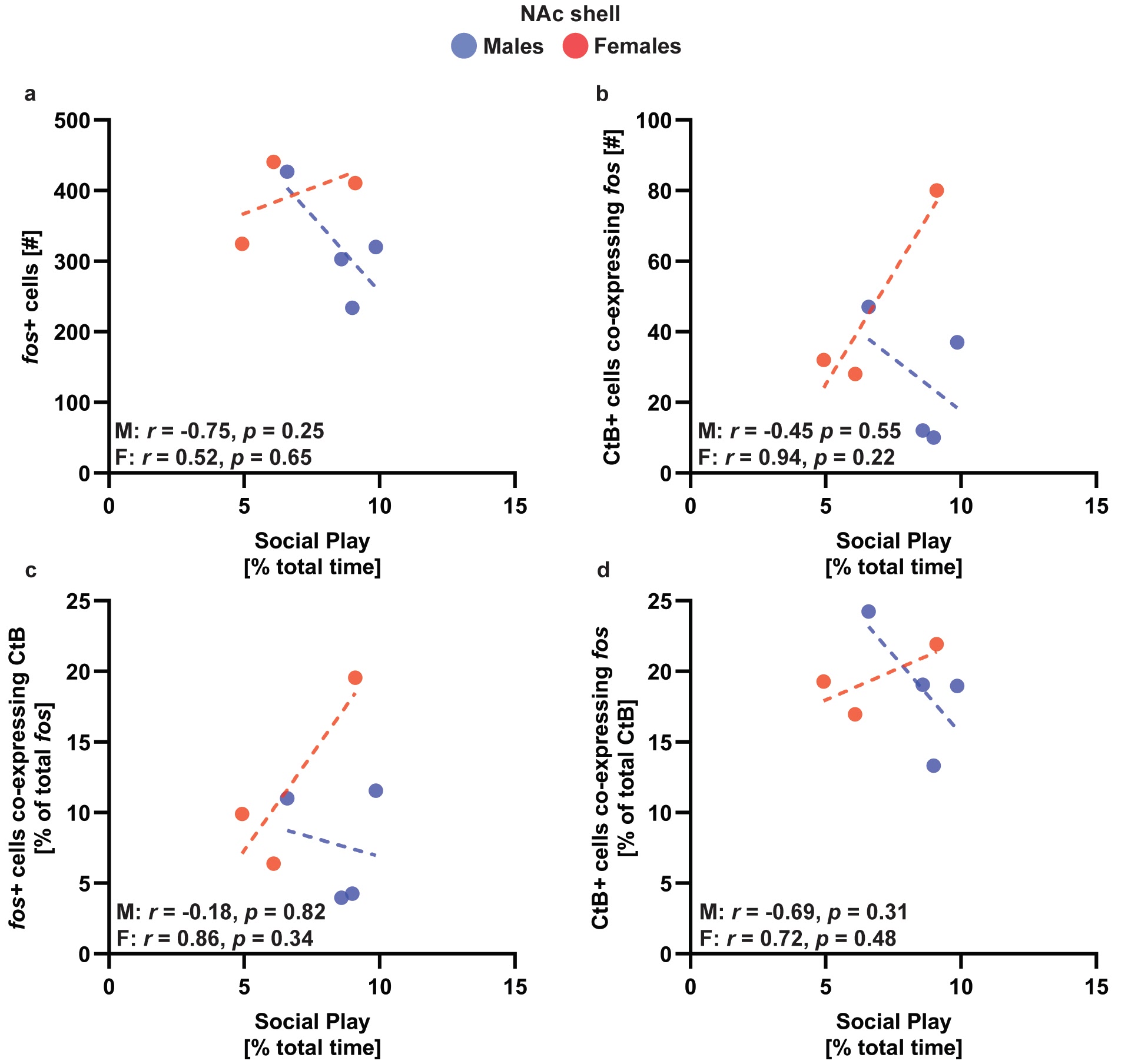

**Supplementary Figure 8. Experiment 3: Social play duration did not significantly correlate with any of the analyzed parameters in the NAc shell.** The percent of time spent engaging in social play did not correlate with the number of *fos*+ cells (a), the number of CtB+ cells co-expressing *fos* (b), the proportion of recruited cells that project to the VP (c), or the proportion of VP-projecting cells activated (d) in the NAc shell of either sex.
